# Leveraging multiple layers of data to predict *Drosophila* complex traits

**DOI:** 10.1101/824896

**Authors:** Fabio Morgante, Wen Huang, Peter Sørensen, Christian Maltecca, Trudy F. C. Mackay

**Author notes:** Section of Genetic Medicine, Department of Medicine, University of Chicago, Chicago, IL 60637, USA. Department of Animal Science, Michigan State University, East Lansing, MI 48824, USA. Center for Human Genetics and Department of Genetics and Biochemistry, Clemson University, Greenwood, SC 29646.

## Abstract

An important challenge in genetics is to be able to predict complex traits accurately. Despite recent advances, prediction accuracy for most complex traits remains low. Here, we used the *Drosophila* Genetic Reference Panel (DGRP), a collection of 200 lines with whole-genome sequences and deep RNA sequencing data, to evaluate the usefulness of using high-quality gene expression levels compared to relying on genotypes for predicting three complex traits. We found that expression levels provided higher accuracy than genotypes for starvation resistance, similar accuracy for chill coma recovery, and lower accuracy for startle response. Models including both genotype and expressions levels did not outperform the best single component model. However, accuracy increased considerably for all the three traits when we included another layer of information, i.e., gene ontology (GO). We found that a limited number of GO terms, some of which had a clear biological interpretation, were strongly predictive of the traits. In summary, this study shows that integrating different sources of information can improve prediction accuracy, especially when large samples are not available.

## Introduction

Predicting complex traits is a fundamental aim of quantitative genetics. Historically, prediction of economically important traits has been important in animal and plant breeding, where the interest lies in predicting breeding values (i.e. the additive genetic part of the phenotype) to select the best individuals for reproduction. Until recently, breeding values were predicted using mixed model methodology and pedigree relationships between individuals (Mrode and Thompson 2005).

However, advances of technology have made genotyping of individuals at several thousands of single nucleotide polymorphisms (SNPs) throughout the entire genome possible and increasingly cheaper. This has led to the possibility of predicting breeding values for traits of interest as a linear combination of SNP genotypes of individuals in a target population and SNP effects estimated in a training population (Meuwissen *et al.* 2001; Goddard and Hayes 2009). The main advantages of this new methodology, termed ‘genomic selection (GS), over the classical phenotypic selection are 1) the higher accuracy of the predicted breeding values due to capturing the Mendelian sampling term; and 2) the shorter generation interval due to the possibility of testing individuals as soon as they are born (Meuwissen *et al.* 2001; Schaeffer 2006; Hayes *et al.* 2009).

GS has been applied to many agricultural species, the first being livestock. Despite the fact that GS has revolutionized animal breeding, the results have varied greatly between species. In dairy cattle, which has the largest and highest quality reference populations, the accuracy of genomic estimated breeding values (gEBVs) has been ∼0.7-0.8 for many traits. This high accuracy combined with the reduced generation interval has doubled the rate of genetic gain since the implementation of GS in breeding programs (Meuwissen *et al.* 2016). However, in beef cattle and pigs, for example, the reference populations are much smaller within each breed, resulting in lower accuracy. To overcome this issue, multi-breed reference populations have been used. However, given the heterogeneous linkage disequilibrium (LD) pattern, across breed prediction has yielded low accuracy (De Roos *et al.* 2009).

One of the most important advantages of GS is that it can be applied to unpedigreed populations since it relies only on actual genotypes. Thus, prediction of complex traits based on genotype data (i.e. genomic prediction, GP) has been used in other branches of genetics, where artificial selection is not the main goal. In evolutionary genetics, the primary interest is in the prediction of evolutionary trajectories of complex traits related to fitness, e.g. body size. In human genetics, we wish to predict the probability of developing a particular disease given an individual’s own genome (Goddard *et al.* 2016).

In human genetics, prediction of complex traits and diseases has been of great interest, especially given the growing effort to implement personalized medicine (de los Campos *et al.* 2010; The Academy of Medical Sciences 2015). Traditionally, human geneticists have used Polygenic Risk Scores (PRS) to predict complex traits. PRS are constructed as a weighted sum of the SNPs associated with the trait of interest from a genome wide association study (GWAS), with the estimated effects used as the weights (Dudbridge 2013; Wray *et al.* 2014). While PRS have had significant predictive power, this has been generally limited (Machiela *et al.* 2011; Ripke *et al.* 2014).

Another approach to GP is by regressing phenotypes on hundreds of variants concurrently using methods borrowed from animal breeding. This class of methods, called whole-genome regression (WGR), has the advantage of accounting for LD among SNPs when estimating their effects (de los Campos *et al.* 2010). However, early attempts at predicting human complex traits using WGR still yielded low prediction accuracy, especially in samples of unrelated individuals (Makowsky *et al.* 2011; de los Campos *et al.* 2013). Recently, using extremely large datasets combined with prediction methods that perform variable selection, much higher prediction accuracy was obtained (Kim *et al.* 2017; Lello *et al.* 2017). These results highlight that adequate sample sizes and methods that account for trait architecture are needed to obtain higher prediction accuracy (Morgante et al. 2018).

In recent years, it has become possible to obtain multiple high quality “omic” data (e.g., gene expression levels, protein levels, metabolite levels) in addition to genotypes for the same samples. This has opened to the possibility of integrating different types of data to uncover genotype-phenotype relationships with a systems genetics approach (Mackay *et al.* 2009; Ritchie *et al.* 2015). The first type of “omic” data to become available on a genome-wide scale was gene expression levels measured using microarray or RNA-seq. One way to use these data is to perform expression quantitative trait loci (eQTL) mapping, to correlate expression levels with genetic variation (Gilad *et al.* 2008). Combining eQTL studies with GWAS revealed that significant hits for many diseases are likely to be eQTLs, meaning that disease SNPs presumably act by altering expression levels (Nicolae *et al.* 2010). As other types of “omic” data become available, studies mapping protein QTL (pQTL) (Chick *et al.* 2016) and metabolic QTL (mQTL) (Kraus *et al.* 2015) have helped elucidate how the effect of genetic variation percolates through intermediate molecular layers before affecting phenotypes.

A complementary way in which multiple “omics” can be used is for complex trait prediction, in a similar manner to prediction using genomic data. Different layers of data may provide (partially) non-redundant information about phenotypes (Guo *et al.* 2016). For example, gene expression levels may also capture some environmental effects, i.e., those that affect levels of expression. In addition, as we move down in the hierarchy genome → epigenome → transcriptome → proteome → metabolome → phenome, the distribution of effects may become simpler, with variables of larger effect that are easier to estimate.

Despite great promise, prediction of complex traits from multiple layers of “omic” data has not been investigated much to date. The first study, to our knowledge, to investigate poly-omic prediction was by Wheeler *et al.* (2014). The researchers developed a method called OmicKriging that was able to incorporate different omics through similarity matrices, one for each “omic”, among individuals. They then applied the method to cell lines with RNA and micro-RNA data, showing that a combined model including both types of “omics” achieved a higher prediction *R*^*2*^ for a quantitative trait than the models including a single type of data. However, when the researchers turned to a clinical dataset that included individuals with both DNA and RNA data and tried to predict a quantitative trait, the combined model predicted worse than the best single component model (Wheeler *et al.* 2014).

Guo *et al.* (2016) used inbred lines of maize with genotype (G), gene expression level (T) and metabolite level (M) information to predict several complex traits using BLUP methodology. In general, MBLUP yielded lower accuracy than all the other models. TBLUP and GBLUP provided similar accuracies, although, on average, GBLUP had better performance. In the majority of cases, poly-omic models (GTBLUP, GMBLUP, GTMBLUP) performed better than single-omic models. However, in many situations, the improvement provided by combined models was minimal (Guo *et al.* 2016).

Vazquez *et al.* (2016) developed a method called Bayesian generalized additive model (BGAM) to incorporate multiple layers of “omic” data, each with a specific prior distribution. Using data from The Cancer Genome Atlas, the researchers showed that integrating multiple “omics” (i.e. gene expression levels, DNA methylation levels and CNV status) generally improved accuracy of prediction of breast cancer survival in humans (Vazquez *et al.* 2016).

Marigorta *et al.* (2017) showed an alternative use of genomic and transcriptomic data. Transcriptional risk scores (TRS) were built using transcript abundance of genes with eQTLs that were in LD with or were inflammatory bowel disease (IBD)-associated SNPs. The researchers then compared the performance of PRS and TRS to distinguish individuals with Crohn’s disease from controls and found that TRS largely outperformed PRS. In addition, TRS were also able to predict disease progression whereas PRS were not (Marigorta *et al.* 2017).

Very recently, Li *et al.* (2019) used genotype, expression (obtained by tiling arrays; Huang *et al.* 2015), and phenotypic data from the *Drosophila melanogaster* Genetic Reference Panel (DGRP) (Mackay *et al.* 2012; Huang *et al.* 2014) to evaluate proportion of variance explained and predictive ability of similar models to Guo *et al.* (2016). The results showed that while models including expression data could capture a greater amount of variance, their predictive ability was generally similar to GBLUP (Li *et al.* 2019).

Although these studies have shown that the use of multiple “omics” to increase the prediction accuracy of complex traits is promising, their results were partially contrasting and, in any case, trait-specific. Thus, in order to elucidate the usefulness of poly-omic prediction of complex traits, we used 200 of the 205 fully sequenced DGRP inbred lines for which gene expression levels were recently obtained by RNA-seq (Everett *et al.* 2019). Taking advantage of the optimal experimental design, high quality “omic” data, and precise phenotype measurements, we sought to evaluate the prediction performance of either single-omic or poly-omic models using three complex traits (starvation resistance, startle response and chill coma recovery) as model traits.

## Materials and Methods

### DGRP lines, genomic, transcriptomic and phenotypic data

The DGRP is a collection of 205 inbred lines derived from 20 generations of full-sib mating from isofemale lines collected at the Farmer’s Market in Raleigh, NC, USA. These lines were fully sequenced using a combination of Illumina and 454 sequencing (Mackay *et al.* 2012; Huang *et al.* 2014). After retaining all the variants with minor allele frequency (MAF) > 0.05 and call rate > 0.8, the total number of variants was 1,891,456.

Recently, an RNA-seq experiment was performed on 200 DGRP lines. Briefly, RNA from whole young adult flies (2-5 days old) raised under standard conditions was extracted and, after depletion of rRNA, sequenced using the Illumina HiSeq 2500 with 125 bp single-end reads. Two biological replicates were obtained for each of the two sexes which were dealt with separately. After all the bioinformatic analyses were performed, line mean expression levels (as log_2_(FPKM)) adjusted for alignment bias for 15,732 and 20,375 genes that were genetically variable across lines in females and males, respectively, were obtained. Of the total number of genes, 11,152 were known genes (i.e. annotated in Flybase) and 4,580 were novel genes in females; on the other hand, 13,249 were known genes and 7,126 were novel genes in males (Everett *et al.* 2019).

To obtain a list of genes that were highly expressed, we pruned out all the genes that had a mean expression across lines < −1.8 log_2_(FPKM). This threshold was obtained from an analysis where a mixture model was fitted to the distribution of all expression values, which was bimodal (Everett *et al.* 2019). This procedure yielded 11,562 and 15,184 highly expressed genes in females and males, respectively.

The DGRP has been phenotyped for many complex traits (Mackay and Huang 2017). Here, we used line means for two fitness traits (starvation resistance and chill coma recovery time) and one behavioral trait (startle response) as model traits (Mackay *et al.* 2012). A total of 198, 172 and 199 lines for starvation resistance, chill coma recovery and startle response, respectively, had both phenotypic measurements and expression levels and were therefore retained for further analyses.

### Statistical analysis – whole genome and transcriptome prediction using linear mixed model

The data were analyzed using several different models. The baseline model was the Genomic Best Linear Unbiased Predictor (GBLUP). This is a linear mixed model where the covariance among lines is modeled using their realized relationships based on DNA marker loci (Habier *et al.* 2007). The model can be written as follows:

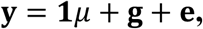

where **y** is an *n*-vector of phenotypes, **1** is an *n*-vector of ones, *μ* is the population mean, **g** is an *n*-vector of random line genomic effects [**g** ∼ *N*(**0, G***σ*^2^_*g*_)] and **e** is an *n*-vector of random residual effects [**e** ∼ *N*(**0, I***σ*^2^_*e*_)]. **G** is the additive genomic relationship matrix (GRM) built using all common variants (MAF > 0.05) according to the formula 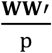 where **W** is the matrix of centered and standardized genotypes for all the lines and p is the number of variants; **I** is the identity matrix.

To evaluate the performance of transcriptomic data for predicting complex traits, we used a Transcriptomic Best Linear Unbiased Predictor (TBLUP). This is very similar to GBLUP, but the GRM is substituted with a transcriptomic relationship matrix (TRM), which evaluates the similarity among lines based on gene expression levels (Guo *et al.* 2016). The model can be written as follows:

**y = 1*μ* + t + e,** where **y** is an *n*-vector of phenotypes, **1** is an *n*-vector of ones, *μ* is the population mean, **t** is an *n*-vector of random line transcriptomic effects [**t** ∼ N(**0, T***σ*^2^_*t*_)] and **e** is an *n*-vector of random residual effects [**e** ∼ N(**0, I***σ*^2^_*e*_)]. **T** is the additive TRM built using only the highly expressed genes, according to the formula 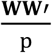 where **W** is the matrix of centered and standardized expression levels for all the lines and p is the number of genes; **I** is the identity matrix.

A combined model, GTBLUP, that had two variance components associated with the GRM and TRM, respectively, was also used. The model can be written as follows:

**y = 1*μ* + g + t + e,** where all the parameters are as defined above.

Finally, a fourth model, GTIBLUP, including three variance components associated with GRM, TRM, and the interaction of the two called IRM, respectively, was fitted. The model can be written as follows:

**y = 1*μ* + g + t + g × t + e,** where **y, 1**, *μ*, **g, t** and **e** are as defined above, and **g**×**t** is an *n*-vector of random line interaction (between genomic and transcriptomic) effects [**g**×**t** ∼ N(**0, G**#**T***σ*^2^_*i*_) where # is the Hadamard product].

The proportion of variance explained in the whole data set by each component was calculated as the ratio of the variance explained by each component over the total phenotypic variance (e.g. in GBLUP, the proportion of variance explained by **g** is equal to 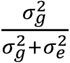).

However, to avoid overfitting in prediction analysis, 30 replicates of 5-fold cross-validation were used; variance components were estimated in the training set, and then used to predict phenotypes in the test set. Prediction accuracy in the test set was evaluated as the squared correlation coefficient, *r*^*2*^, between true and predicted phenotypes, averaged over folds and replicates.

### Statistical analysis – whole genome and transcriptome prediction using Random Forest

Hypothesizing that genes may affect traits at least partially through non-linear interactions, a completely non-parametric method, the Random Forest (Breiman 2001), was used to potentially capture those interaction effects. The algorithm implemented in the ‘randomForest’ R package was fitted using the expression levels of highly expressed genes as predictors, with default values of the tuning parameters and 1,000 trees. This analysis was performed using the same cross-validation scheme and metrics as the whole genome and transcriptome analysis.

### Statistical analysis – transcriptome-wide association study (TWAS) informed prediction

To try and enrich the TBLUP model for genes associated with the trait of interest and to eliminate noise from unassociated genes, variable selection was performed re-adapting the approach of Morgante et al. (2018). At each round of cross-validation, TWAS, i.e. regressing the phenotype on the expression level of each gene one at a time, was performed in the training set. Only highly expressed genes were used. The genes with *P* < X (X = 5*10^−1^; 10^−1^; 10^−2^; 10^−3^; 10^−4^; 10^−5^; 10^−6^) for their main effect were selected and used to build a trait-specific TRM. This trait-specific TRM was fitted in the TBLUP model to estimate variance components in the training set to be used to predict phenotypes in the test set.

As a control, prediction using only randomly sampled genes was performed for the three traits. At each round of cross-validation, k genes (k = 5; 50; 500; 1,000; 5,000) were randomly sampled from the set of highly expressed genes and used for prediction in a similar way to the TWAS-selected genes.

### Statistical analysis – gene ontology (GO) informed prediction

With the same aim to disentangle signal from noise, we used a different procedure that relies on external sources of information, e.g. functional annotation. Edwards *et al.* (2016) showed that exploiting information about gene ontology categories could improve the accuracy of SNP-based prediction. Here, we followed the same approach for SNP-based prediction with our data and extended it to expression-based prediction. SNPs were mapped to genes based on FlyBase v. 5.57 annotation (St Pierre *et al.* 2014). Genes were then mapped to GO terms using the R package ‘org.Dm.eg.db’ v. 3.4.0 (Carlson 2016) available in BioConductor.

For SNP-based prediction, the following model (GO-GBLUP) was fitted:

**y = 1*μ* + g**_**GO**_ **+ g**_**notGO**_ **+ e,** where **g**_**GO**_ is an *n*-vector of random line genomic effects associated with SNPs pertaining to a specific GO term (through a GO-specific GRM built using SNPs in a specific GO), **g**_**notGO**_ is an *n*-vector of random line genomic effects associated with all the remaining SNPs (through a GRM built using all SNPs not in that GO), and all the other parameters are as defined above. This model was fitted for all GO terms including at least 5 genes, resulting in 2,306 GO terms.

For expression-based prediction, the following model (GO-TBLUP) was fitted: **y = 1*μ* + t**_**GO**_ **+ t**_**notGO**_ **+ e,** where **t**_**GO**_ is an *n*-vector of random line transcriptomic effects associated with genes pertaining to a specific GO term (through a GO-specific TRM built using genes in a specific GO), **t**_**notGO**_ is an *n*-vector of random line transcriptomic effects associated with all the remaining genes (through a TRM built using all genes not in that GO), and all the other parameters are as defined above. This model was fitted for all GO terms including at least 5 genes and that were present in our expression data restricted to highly expressed genes, resulting in 2,089 and 2.046 GO terms for females and males, respectively.

In order to evaluate whether genome-level and transcriptome-level GO terms contribute overlapping information, the following model (GO-GTBLUP) was fitted: **y = 1*μ* + t**_**GO**_ **+ t**_**notGO**_ **+ g**_**GO**_ **+ g**_**notGO**_ **+ e,** where all the parameters are as defined above. This model was fitted for all GO terms that were in common between GO-SNPs and GO-genes after pruning according to the requirements described above, resulting 2,083 and 2.039 GO terms for females and males, respectively.

All these analyses were performed using the same cross-validation scheme and metrics as the whole genome and transcriptome analysis.

## Results

Throughout this manuscript, the results of all the analyses for starvation resistance are presented in the main text, while the results for startle response and chill coma recovery are presented in the supplementary material.

### Whole genome and transcriptome prediction using linear mixed models

The first step of the analysis was to determine to what extent the transcriptome could explain the phenotypic variance in the whole data set for the traits of interest, compared to the variance explained by the genome. To do that, we fitted TBLUP and GBLUP to the whole data set. Figure 1 shows that both GBLUP and TBLUP were able to explain the large majority or even all of the phenotypic variability for starvation resistance in females and males. However, while GBLUP was able to explain about 50% of the total phenotypic variability for startle response, TBLUP could only explain about 25% of the trait variation for females and males (Fig. S1). For chill coma recovery in females, GBLUP was able to explain the majority of variance, but TBLUP was able to explain only minimal variance. However, neither model was able to explain any phenotypic variance in males (Fig. S2).

**Figure 1.**
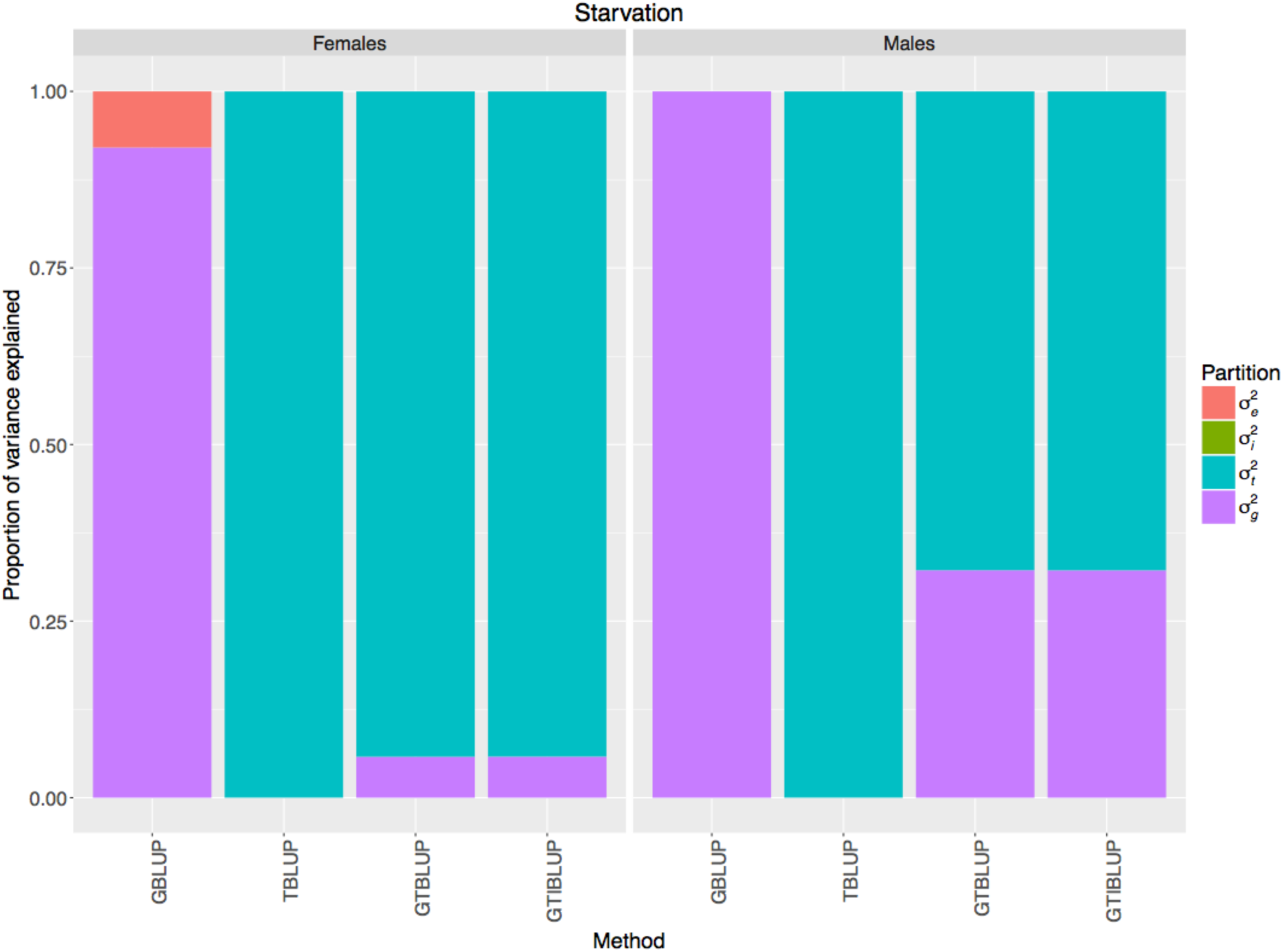
Proportion of phenotypic variance for starvation resistance explained by each component in the models fitted to the whole data set, i.e. GBLUP (*σ*^2^_*g*_ and *σ*^2^_*e*_), TBLUP (*σ*^2^_*t*_ and *σ*^2^_*e*_), GTBLUP (*σ*^2^_*g*_, *σ*^2^_*t*_ and *σ*^2^_*e*_), and GTIBLUP (*σ*^2^_*g*_, *σ*^2^_*t*_, *σ*^2^_*i*_ and *σ*^2^_*e*_). The left panel represents females, and the right panel represents males.

We then fitted GTBLUP to evaluate the relative contribution of the genome and the transcriptome to explain variance. This model was able to explain all the phenotypic variance for starvation resistance in females and males, with the transcriptome contributing much more than the genome (Fig. 1). For startle response, the proportion of variance explained by GTBLUP was very similar to that explained by GBLUP, with the genome contributing much more variance than the transcriptome in both females and males (Fig. S1). For chill coma recovery, GTBLUP could explain roughly the same amount of variance as GBLUP in females, almost totally driven by the variance explained by the genome. On the other hand, in males, no variance could be explained by GTBLUP (Fig. S2).

The results of GTIBLUP showed that the interaction between genome and transcriptome did not explain any phenotypic variance, except for startle response and chill coma recovery in females (Figs. 1, S1, S2).

Having shown that the transcriptome can capture some signal in the majority of cases, we proceeded by evaluating its predictive ability in a cross-validation setting. For starvation resistance and startle response, prediction accuracy in the test set showed the same patterns as variance explained in the whole data, although much lower. For starvation resistance, the highest prediction accuracy was given by TBLUP, which was roughly twice as high as that from GBLUP; GTBLUP and GTIBLUP did not improve accuracy over TBLUP (Fig. 2). For startle response, GBLUP provided the highest prediction accuracy, which was roughly twice as high as that from TBLUP; GTBLUP and GTIBLUP had accuracies about half way between GBLUP and TBLUP (Fig. S3). On the other hand, for chill coma recovery all the four models provided more similar prediction accuracies, which were lower and much more variable (i.e. with larger standard error) than for the other two traits (Fig. S4).

**Figure 2.**
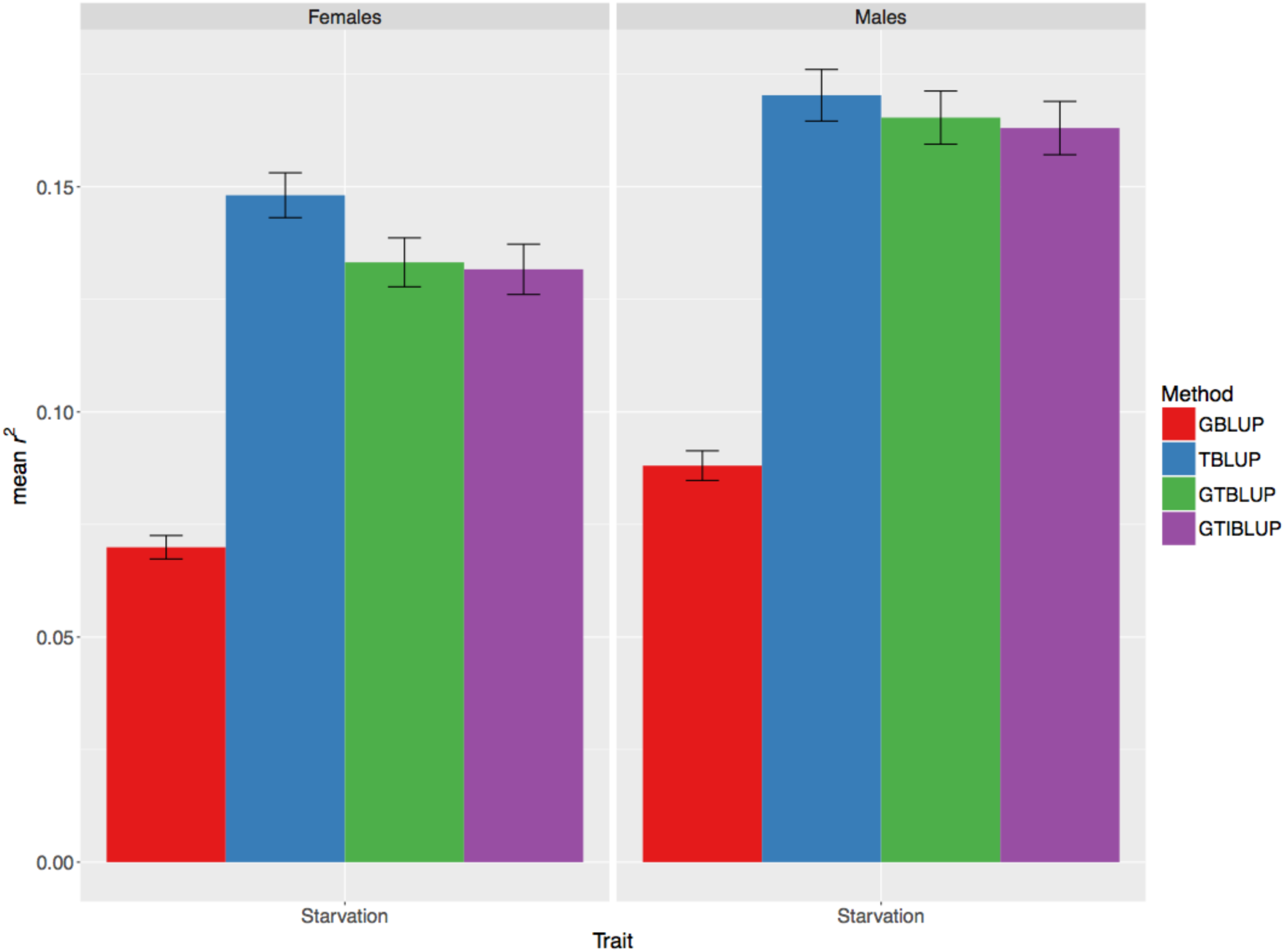
Prediction accuracy for starvation resistance, measured as mean *r*^*2*^ (on the y-axis) in the test set, obtained by GBLUP, TBLUP, GTBLUP, and GTIBLUP. The bars represent the standard error of the mean. The left panel represents females, and the right panel represents males.

### Whole genome and transcriptome prediction using Random Forest

To assess whether non-parametric methods could perform better than linear models by capturing some potential interactions among genes, the Random Forest was fitted to the data. The results showed that the Random Forest did not perform consistently better than TBLUP for any trait (Figs. 3, S5, S6).

**Figure 3.**
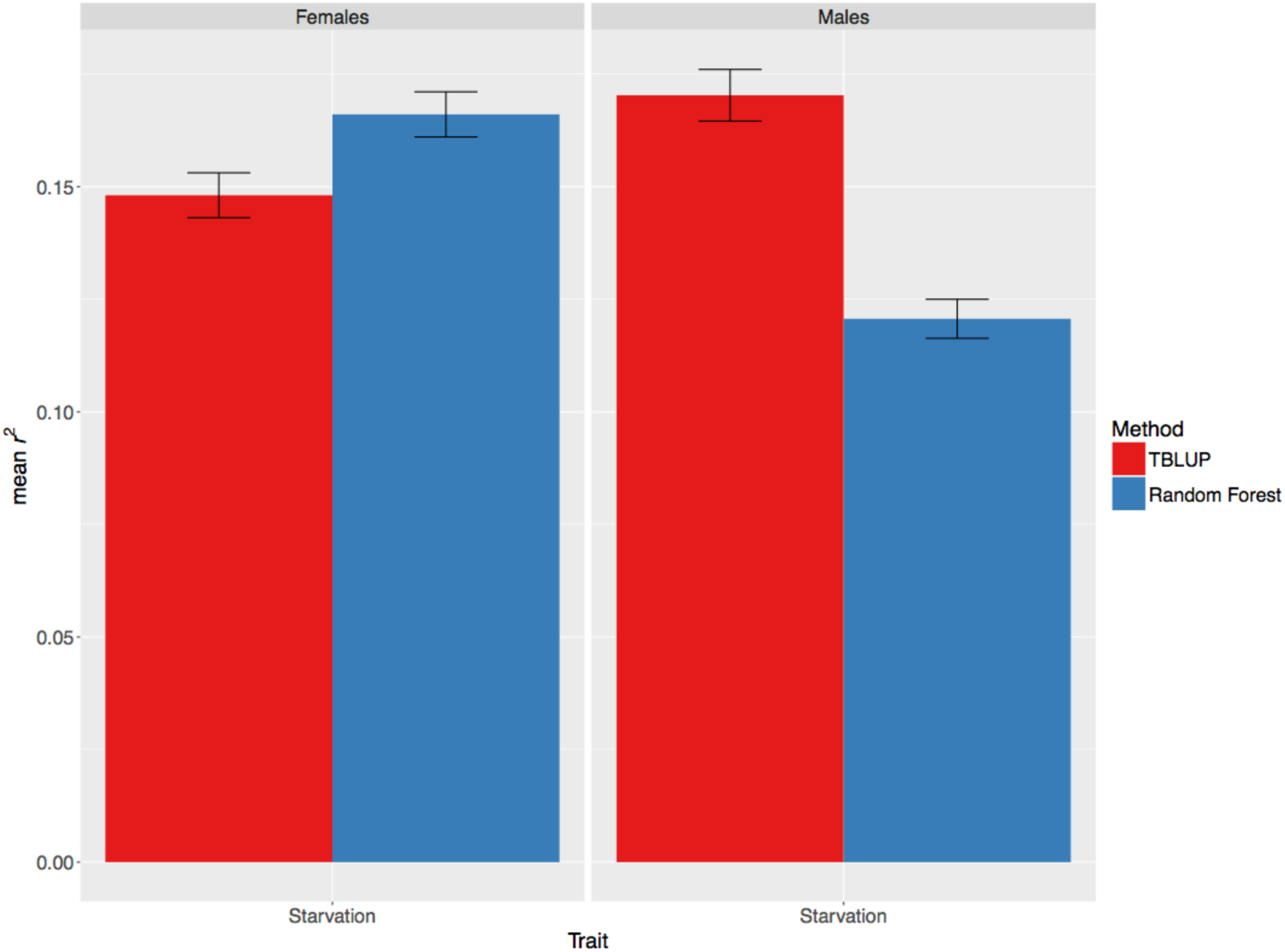
Comparison between the prediction accuracy for starvation resistance, measured as mean *r*^*2*^ (on the y-axis) in the test set, obtained by TBLUP and Random Forest. The bars represent the standard error of the mean. The left panel represents females, and the right panel represents males.

### Transcriptome-wide association study (TWAS) informed prediction

We combined mapping and prediction into a single pipeline in order to enrich the TBLUP model for genes associated with the trait of interest. The results of this analysis revealed no specific relationship between prediction *r*^*2*^ and p-value threshold for the trait/sex combinations (Figs. 4, S7, S8). For example, for starvation resistance in females, the most stringent p-value threshold (i.e. 10^−6^) improved prediction accuracy over using all highly expressed genes and gave the best accuracy overall. However, the same threshold yielded among the worst accuracies for starvation resistance in males (Fig. 4). On the other hand, for startle response in males, the most lenient threshold (*P* < 5*10^−1^) doubled the accuracy given by all highly expressed genes while more stringent thresholds yielded lower accuracy (Fig. S7).

**Figure 4.**
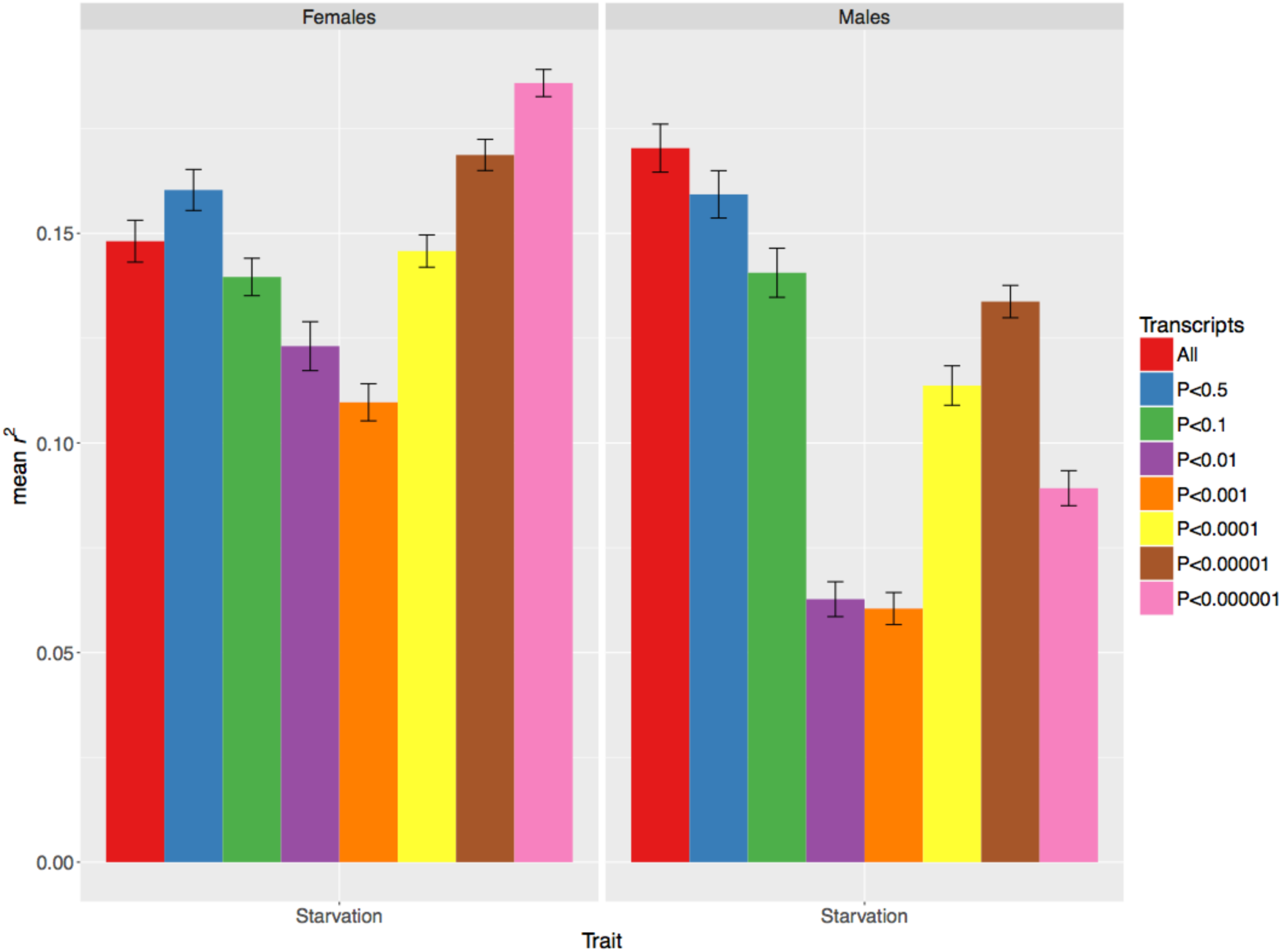
Prediction accuracy for starvation resistance, measured as mean *r*^*2*^ (on the y-axis) in the test set, obtained by the TWAS-TBLUP model at different p-value thresholds for selecting genes. The bars represent the standard error of the mean. The left panel represents females, and the right panel represents males.

The results of the analysis using only randomly sampled genes showed that only a few genes were able to give very similar accuracies as those given by the whole transcriptome (Figs. 5, S9, S10). This trend was more pronounced for startle response and chill coma recovery (Figs. S9, S10); this observation might be due to the much lower accuracy obtained with all highly expressed genes to begin with.

**Figure 5.**
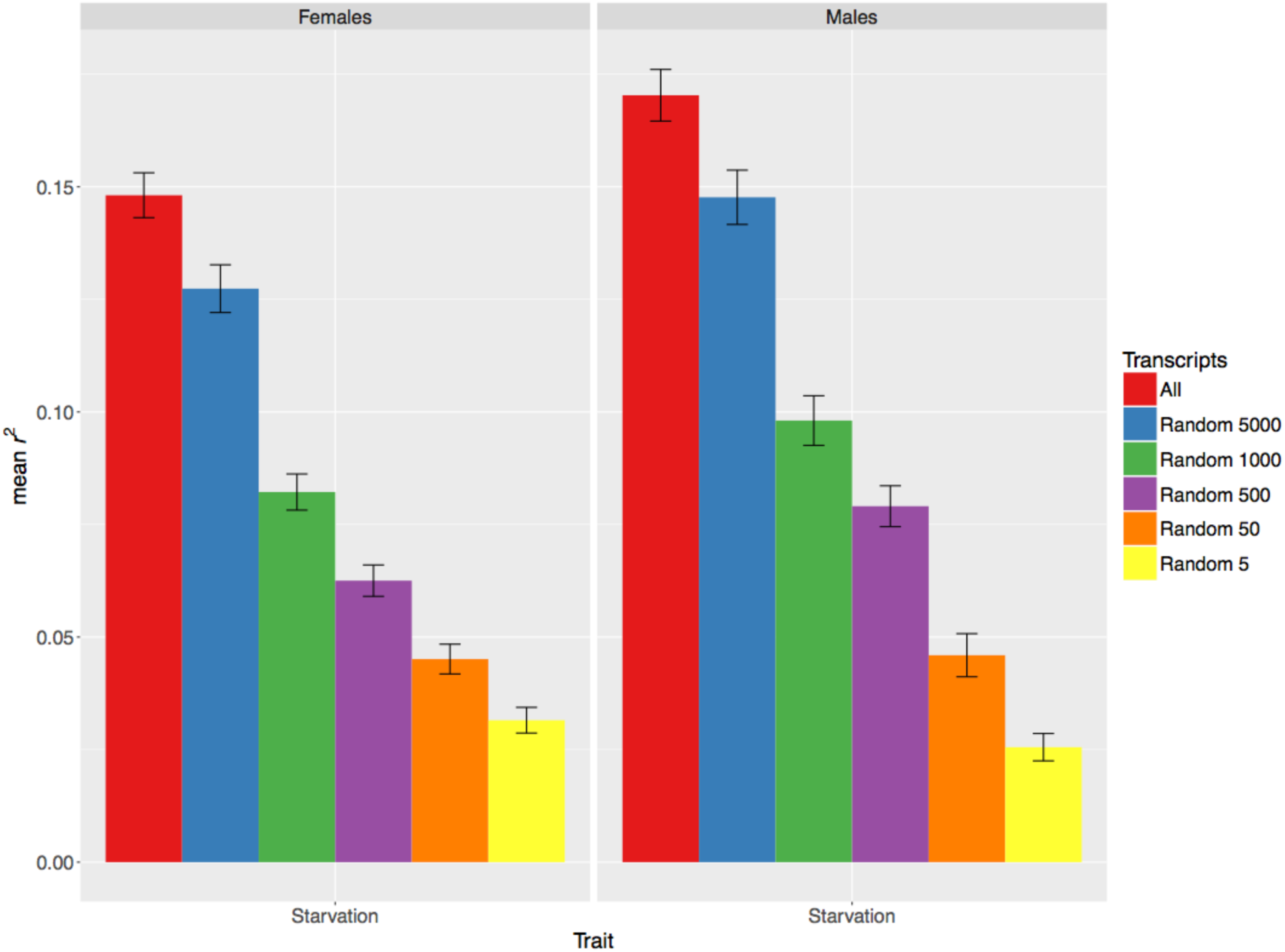
Prediction accuracy for starvation resistance, measured as mean *r*^*2*^ (on the y-axis) in the test set, obtained by using only different numbers of randomly selected genes in the TBLUP model. The bars represent the standard error of the mean. The left panel represents females, and the right panel represents males.

Comparing the two sets of results, it was obvious that the TWAS-TBLUP strategy was not effective for the three traits analyzed.

### Gene ontology (GO) informed prediction

We hypothesized that the lack of success of the TWAS-TBLUP model might have been due to the small sample size of the DGRP, which could make mapping difficult. Thus, to disentangle signal from noise, we adopted a procedure that relies on functional annotation to group SNPs and genes according to GO categories, and check whether some GO terms are particularly predictive of the traits.

First, we fitted the same GO-GBLUP model that was proposed by Edwards *et al.* (2016). The results showed that while the majority of GO terms provided similar accuracy to the baseline GBLUP model (the black horizontal line in the graphs), some GO terms achieved much higher *r*^*2*^ values. This pattern was observed for all the trait/sex combinations (Figs. 6, S11, S12). Some of the most predictive GO terms had a clear interpretation. For example, the most predictive GO term for starvation resistance in males, GO:0034389, has been implicated in lipid particle organization according to FlyBase (Gramates *et al.* 2017).

**Figure 6.**
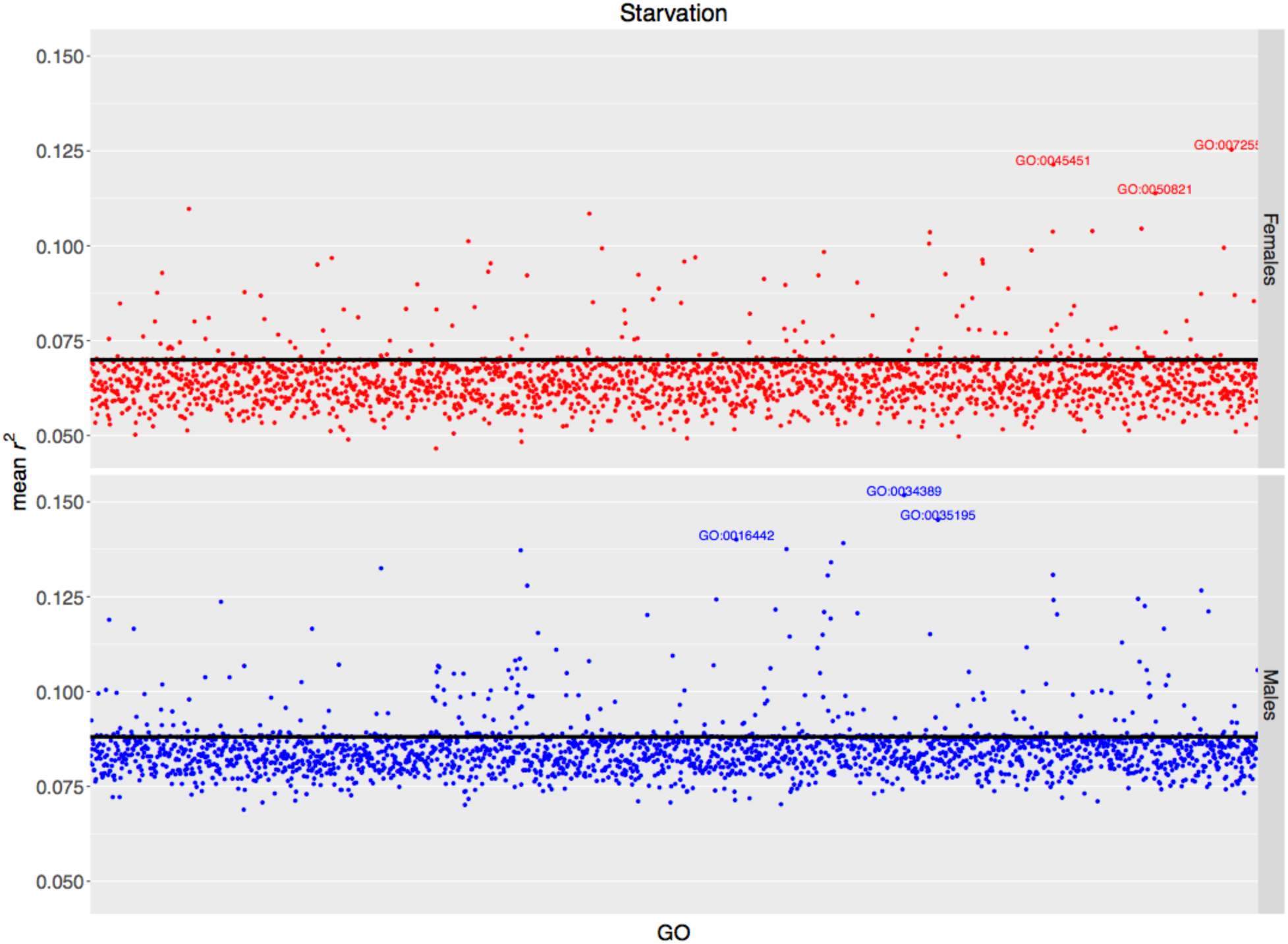
Prediction accuracy for starvation resistance, measured as mean *r*^*2*^ (on the y-axis) in the test set, obtained by the GO-GBLUP model. Each point represents the prediction accuracy achieved by a specific GO term; the top 3 most predictive GO terms are spelled out. The black horizontal line represents the accuracy of the baseline GBLUP model. The upper panel represents females, and the lower panel represents males.

Then, using the same rationale, we developed the GO-TBLUP model and fitted it to our data. The results showed a very similar pattern to those of GO-GBLUP – most GO terms had very similar *r*^*2*^ values to TBLUP (the black horizontal line in the graphs), yet some GO terms provided much higher accuracies (Figs. 7, S13, S14). Again, some of the most predictive GO terms had a clear interpretation. For example, the most predictive GO term for starvation resistance in females, GO:0033500, has been implicated in carbohydrate homeostasis according to FlyBase (Gramates *et al.* 2017). Interestingly, the most predictive SNP-based GO terms and the most predictive gene-based GO terms were distinct.

**Figure 7.**
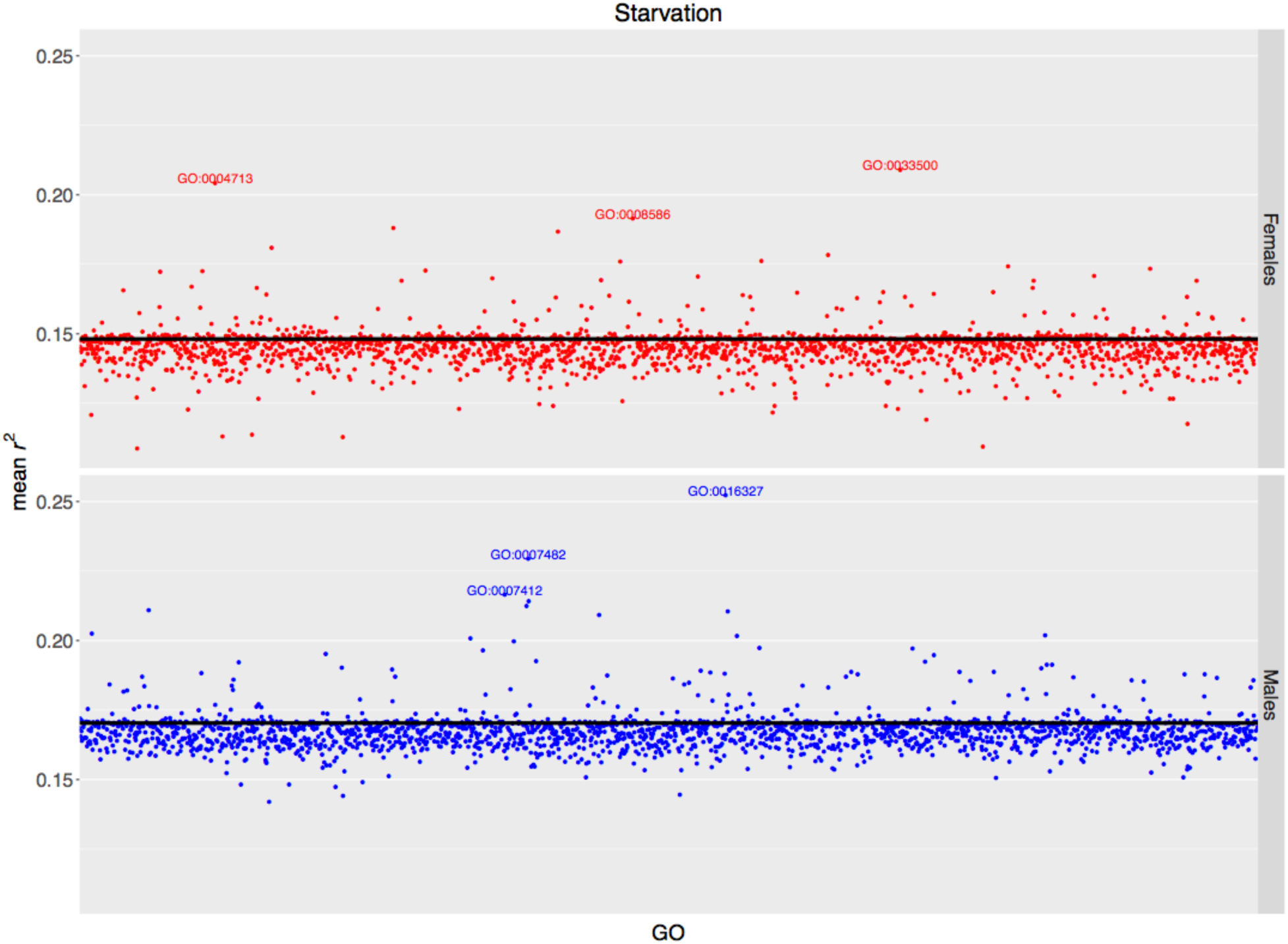
Prediction accuracy for starvation resistance, measured as mean *r*^*2*^ (on the y-axis) in the test set, obtained by the GO-TBLUP model. Each point represents the prediction accuracy achieved by a specific GO term; the top 3 most predictive GO terms are spelled out. The black horizontal line represents the accuracy of the baseline TBLUP model. The upper panel represents females, and the lower panel represents males.

Lastly, we fitted a GO-GTBLUP model to see whether genomic data and transcriptomic data contributed overlapping information to prediction accuracy for each GO. Generally, the combined model did not yield higher accuracy than the best between GO-GBLUP and GO-GBLUP; however, the ranking of the most predictive GO terms might have changed slightly (Figs. 8, S15, S16). Only in one case, chill coma recovery in males, did this combined model achieve the best accuracy, although only marginally (Fig. S16).

**Figure 8.**
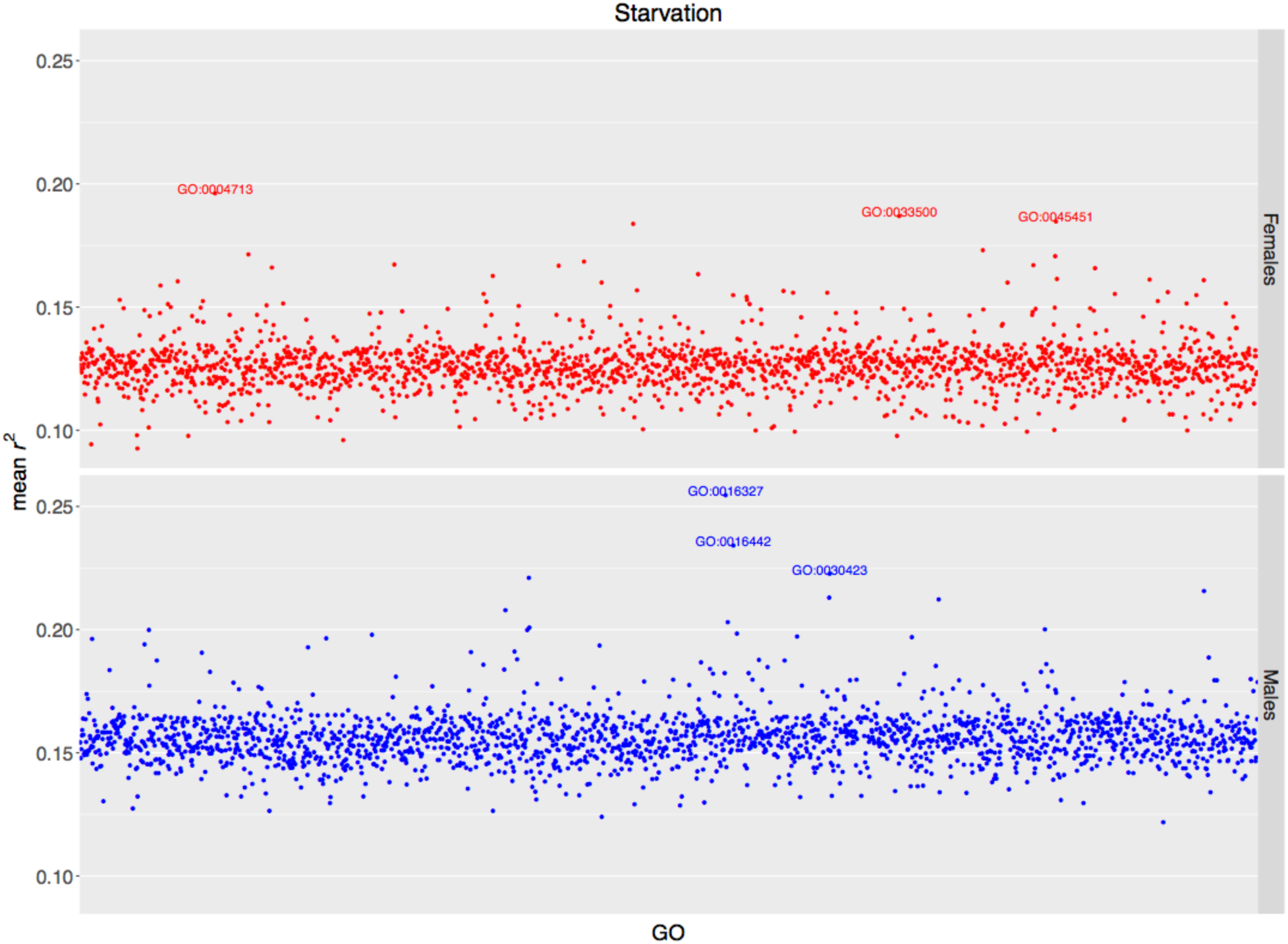
Prediction accuracy for starvation resistance, measured as mean *r*^*2*^ (on the y-axis) in the test set, obtained by the GO-GTBLUP model. Each point represents the prediction accuracy achieved by a specific GO term; the top 3 most predictive GO terms are spelled out. The upper panel represents females, and the lower panel represents males.

## Discussion

Here, we evaluated the use of transcriptomic data for the prediction of complex traits, taking advantage of the unique resource of the DGRP, for which RNA-sequence data has been obtained recently (Everett *et al.* 2019).

To assess whether genetic variation in gene expression among lines could explain any phenotypic variability in our data, we fitted TBLUP to the whole data set. We found that the transcriptome could explain at least some phenotypic variance for all the traits except for chill coma recovery in males. When compared with the variance explained by the genome in GBLUP, we found that both the genome and transcriptome could explain about the same proportion of variance for starvation resistance. However, the genome was able to explain a higher proportion of variance for startle response and chill coma recovery. When fitting GTBLUP, we observed that this combined model was generally not able to explain a significantly larger proportion of variance than the best of the GBLUP and TBLUP models. The component that explained the majority of the variance in GTBLUP was the one that performed better in the single-component models, e.g. the transcriptome for starvation resistance. The results of GTIBLUP highlighted the fact that the interaction term contributed to explaining variance only for startle response and chill coma recovery in females. Again, the total proportion of variance explained by GTIBLUP was only marginally higher than GBLUP and GTBLUP for these two trait/sex combinations. Overall, these results seem to suggest that genome and transcriptome may contribute largely overlapping information, with the key player being dependent on the trait analyzed. This contrasts with the results of Guo *et al.* (2016) and Li *et al.* (2019) where the genome consistently explained more variance than the transcriptome, and, in Guo *et al.* (2016), the interaction term contributed to explaining variance for almost all of traits analyzed.

The general trend observed for phenotypic variance explained in the whole data set was confirmed by the results for prediction accuracy in the cross-validation setting. Prediction accuracy was low to very low for all the traits, with a large gap between the proportion of variance explained and prediction *r*^*2*^, confirming what has been observed elsewhere with genomic data (Morgante et al. 2018). The only exception was chill coma recovery in males, for which no model could explain any variance in the whole data, yet some very low but significant prediction accuracy was obtained in the test set with the same models. This seemingly inconsistent observation can be explained by the fact that some variance could be explained in some training-test partitions in the cross-validation procedure, resulting in non-zero prediction accuracy overall.

In general, chill coma recovery had lower and more variable (i.e. larger error standard error) prediction accuracy than the other two traits with any of the models used. This might be due to a more complex genetic architecture of this trait, with a large number of epistatic interactions (Huang *et al.* 2012; Ober *et al.* 2015). For starvation resistance, TBLUP provided the highest accuracy; however, GBLUP provided the highest accuracy for startle response. GTBLUP and GTIBLUP were never better than the best single-component only, adding evidence to the hypothesis that genome and transcriptome contribute largely redundant information.

Our results agreed with those of Guo *et al.* (2016) regarding the absence of an improvement in predictive ability when including the genome-transcriptome interaction term in the model. However, while our results also showed no improvement when using GTBLUP, Guo *et al.* (2016) observed an improvement with the same combined model. One reason for this discrepancy could be that we utilized only genes whose expression levels were genetically variable across lines. This filter was necessary because the individual flies that were phenotyped were not the same flies from which RNA was extracted, despite being of identical genotypes. Thus, genetics was the only link between these two sets of flies. However, this implied that we might have missed some information from genes whose expression levels were not genetically variable but may have captured some (micro-) environmental effects. Our results also differed from those in Li *et al.* (2019) in that they found TBLUP to be consistely much worse than GBLUP, providing null prediction accuracy for many traits. One hypothesis for this observation is the much noisier expression measurments from tiling arrays compared to RNA-seq (Everett *et al.* 2019).

In our previous work, we showed that variable selection was the key to achieve high genomic prediction accuracy (Morgante et al. 2018). Thus, we applied the same strategy to gene expression data. However, the results did not show a clear advantage in using this procedure and were very variable across traits and p-value threshold. There are at least two possible reasons for this observation. First, the DGRP has a small sample size, which can be problematic for mapping genes with smaller effects. Second, expression levels displayed a strong correlation structure across genes (Everett *et al.* 2019); this was highlighted by the fact that a few randomly sampled genes could achieve very similar accuracies to all highly expressed genes. Correlations among genes can be an issue for mapping; correlations may induce spurious associations if they result from some type of structure (e.g. spatial or temporal) present in the expression data. This concept is closely related to population structure with genomic data (Astle and Balding 2009).

To overcome the issues associated with mapping, we sought to enrich our models for genes associated with the traits by exploiting external information. Following Edwards *et al.* (2016), we grouped SNPs into genes and then into GO terms. In agreement with Edwards *et al.* (2016), we found that a limited number of GO terms provided much higher accuracy with GO-GBLUP than the baseline GBLUP. However, although both our work and that of Edwards *et al* (2016) used the DGRP, there were some differences. First, the two studies used different subsets of lines, which can make a difference with such a small sample size. Second, we used 5-fold cross-validation while they used 10-fold cross-validation; the size of the training set has been shown to influence prediction accuracy (Ober *et al.* 2012). Third, we used line means for phenotypes, whereas they used individual measurements. Fourth, we used GO terms containing at least 5 genes while Edwards *et al* (2016) used GO terms containing at least 10 genes. This resulted in the most predictive GO terms being different between the two studies but the general ranking being pretty consistent.

We then extended the methodology of Edwards *et al.* (2016) to work with gene expression directly: GO-TBLUP. We found a similar pattern to the GO-GBLUP results, whereby a small number of GO terms achieved a much higher accuracy than the baseline TBLUP. However, the most predictive GO terms in GO-GBLUP and the most predictive GO terms in GO-TBLUP were not the same. This suggests the possibility that the genome and transcriptome as a whole may contribute redundant information, but they may not when gene ontology information is incorporated. This observation has important implications because it may be possible to build a trait-specific model with the most predictive SNP-based GO terms and the most predictive gene-based GO terms to improve the overall prediction accuracy. However, to be able to do so, it is necessary to develop a procedure to select the most predictive GO terms without bias in the training set, and more research is needed in that area.

In summary, this study has confirmed that using transcriptomic data to predict is promising. Our work, together with other studies (Finucane *et al.* 2015; Edwards *et al.* 2016; Abdollahi-Arpanahi *et al.* 2017), has shown that integrating multiple layers of “omic” data together with functional annotation can identify features that are important to understand and predict complex traits.

This study has some limitations, in addition to using only genes with genetically variable expression levels. First, the DGRP has very small sample size; this limits the maximum accuracy reachable. Second, RNA was extracted from whole flies; this approach gives the average gene expression levels across all tissues. This may be advantageous, because identifying tissues relevant to specific traits may not be easy; however, tissues that were not relevant to the traits analyzed added noise to the expression levels, potentially affecting prediction accuracy. In addition, tissue specificity of gene expression has been demonstrated in human studies (Aguet *et al.* 2017). Third, RNA was extracted from flies that were reared in standard conditions and were not subjected to any external stimulus; however, all three traits analyzed were stress-based. This might be one reason for the poor prediction accuracy of TBLUP for startle response and chill coma recovery. However, for starvation resistance, expression levels on baseline flies may reflect their ability to store and elaborate energetic resources. This might explain the higher accuracy of TBLUP and agrees with the most predictive GO term being implicated in carbohydrate homeostasis.

## Data availability

The genetic and phenotypic data used in this analysis can be found at http://dgrp2.gnets.ncsu.edu. The gene expression data can be found in Everett *et al.* (2019).

## Acknowledgements

This work was supported by Genomic Selection in Animals and Plants (GenSAP) funded by The Danish Council for Strategic Research to T. F. C. M., P. S. and F. M.

## Author contributions

F. M., W. H., P. S., C. M. and T. F. C. M. designed the experiment. F. M. performed the analyses. F. M. and T. F. C. M. wrote the manuscript.

## Author information

The authors declare that no competing interests exist. Correspondence and requests for materials should be addressed to

## Supplementary Material

**Figure S1.**
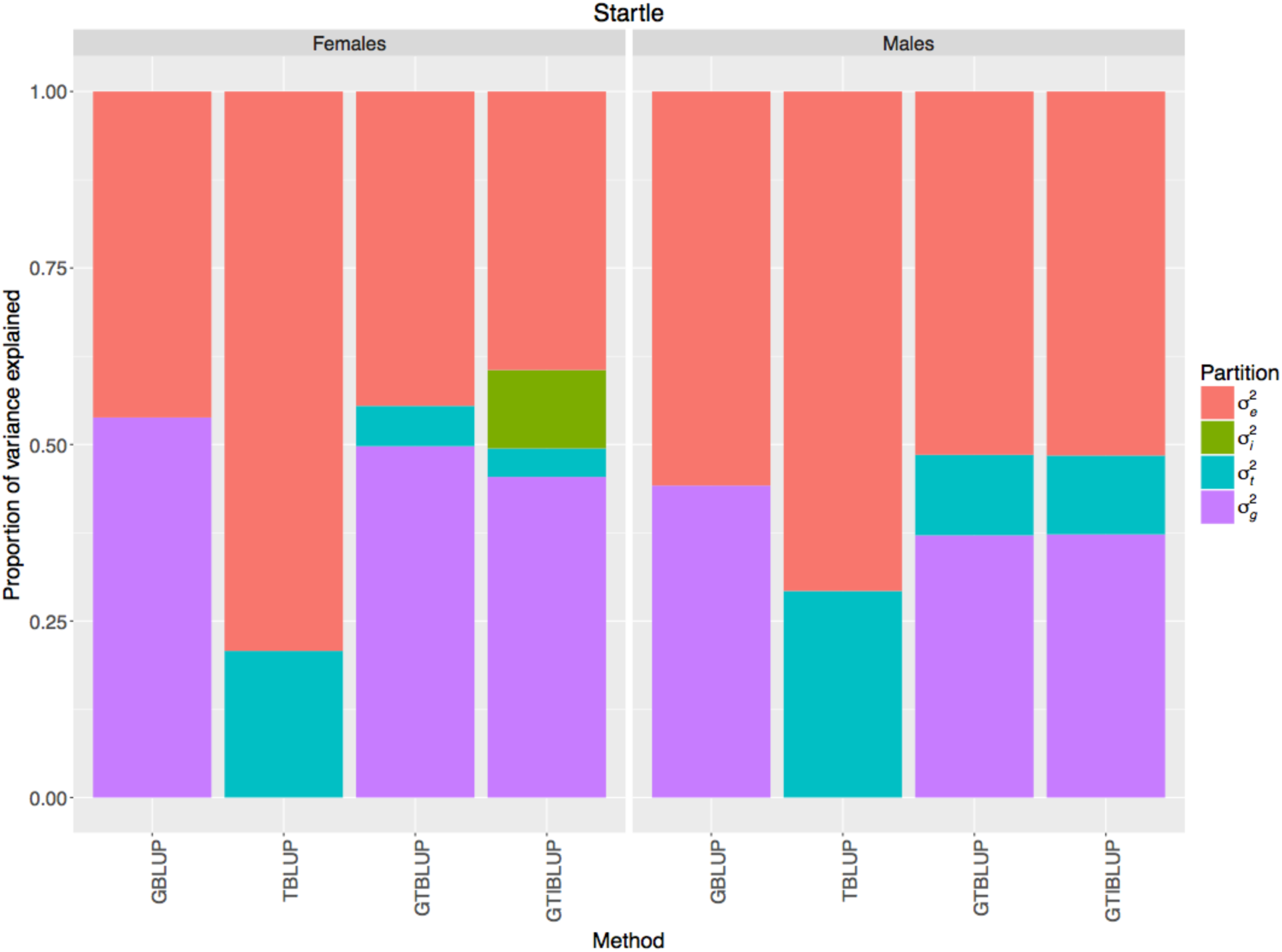
Proportion of phenotypic variance for startle response explained by each component in the models fitted to the whole data set, i.e. GBLUP (*σ*^2^_*g*_ and *σ*^2^_*e*_), TBLUP (*σ*^2^_*t*_ and *σ*^2^_*e*_), GTBLUP (*σ*^2^_*g*_, *σ*^2^_*t*_ and *σ*^2^_*e*_), and GTIBLUP (*σ*^2^_*g*_, *σ*^2^_*t*_, *σ*^2^_*i*_ and *σ*^2^_*e*_). The left panel represents females, and the right panel represents males.

**Figure S2.**
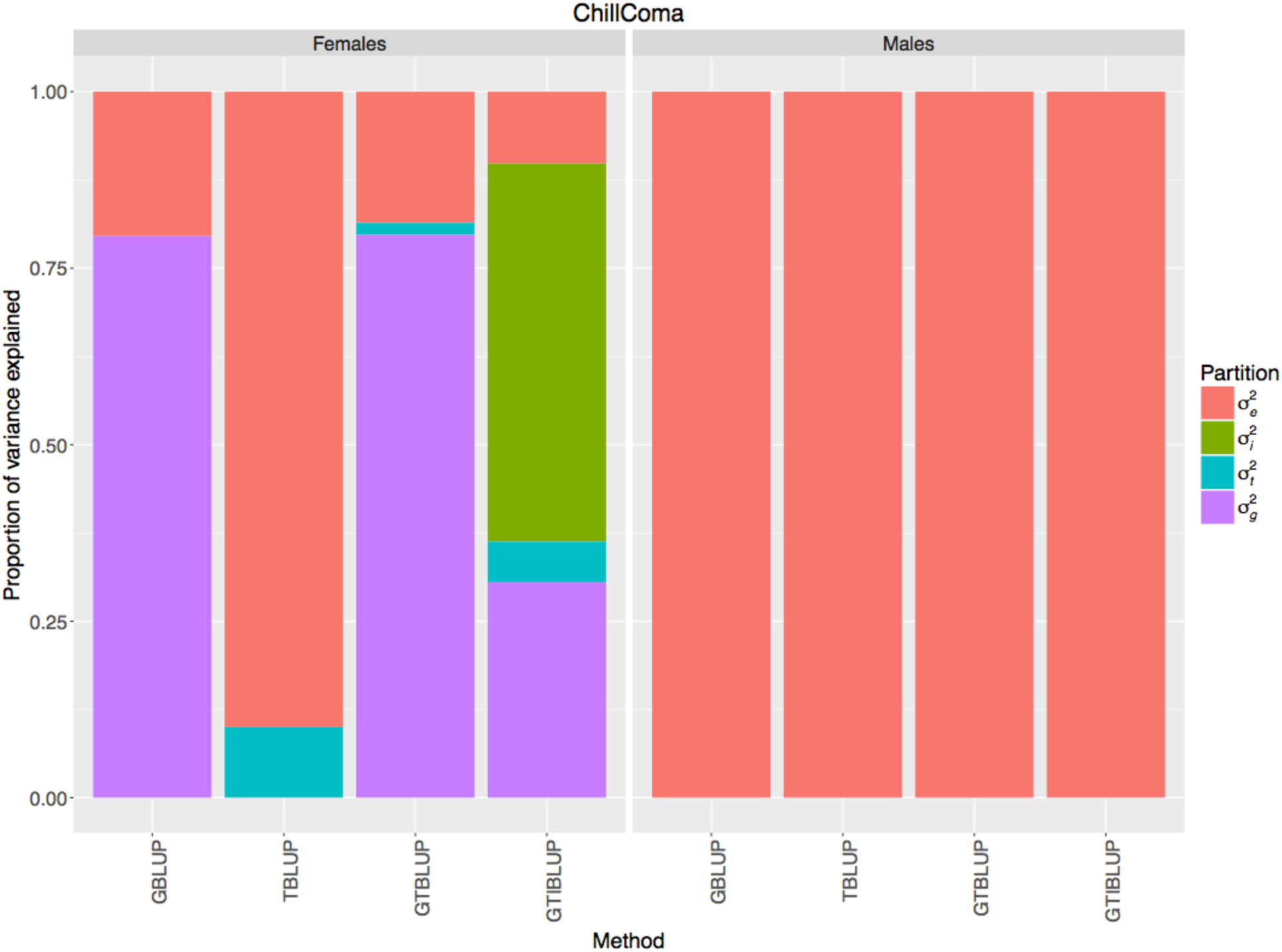
Proportion of phenotypic variance for chill coma recovery explained by each component in the models fitted to the whole data set, i.e. GBLUP (*σ*^2^_*g*_ and *σ*^2^_*e*_), TBLUP (*σ*^2^_*t*_ and *σ*^2^_*e*_), GTBLUP (*σ*^2^_*g*_, *σ*^2^_*t*_ and *σ*^2^_*e*_), and GTIBLUP (*σ*^2^_*g*_, *σ*^2^_*t*_, *σ*^2^_*i*_ and *σ*^2^_*e*_). The left panel represents females, and the right panel represents males.

**Figure S3.**
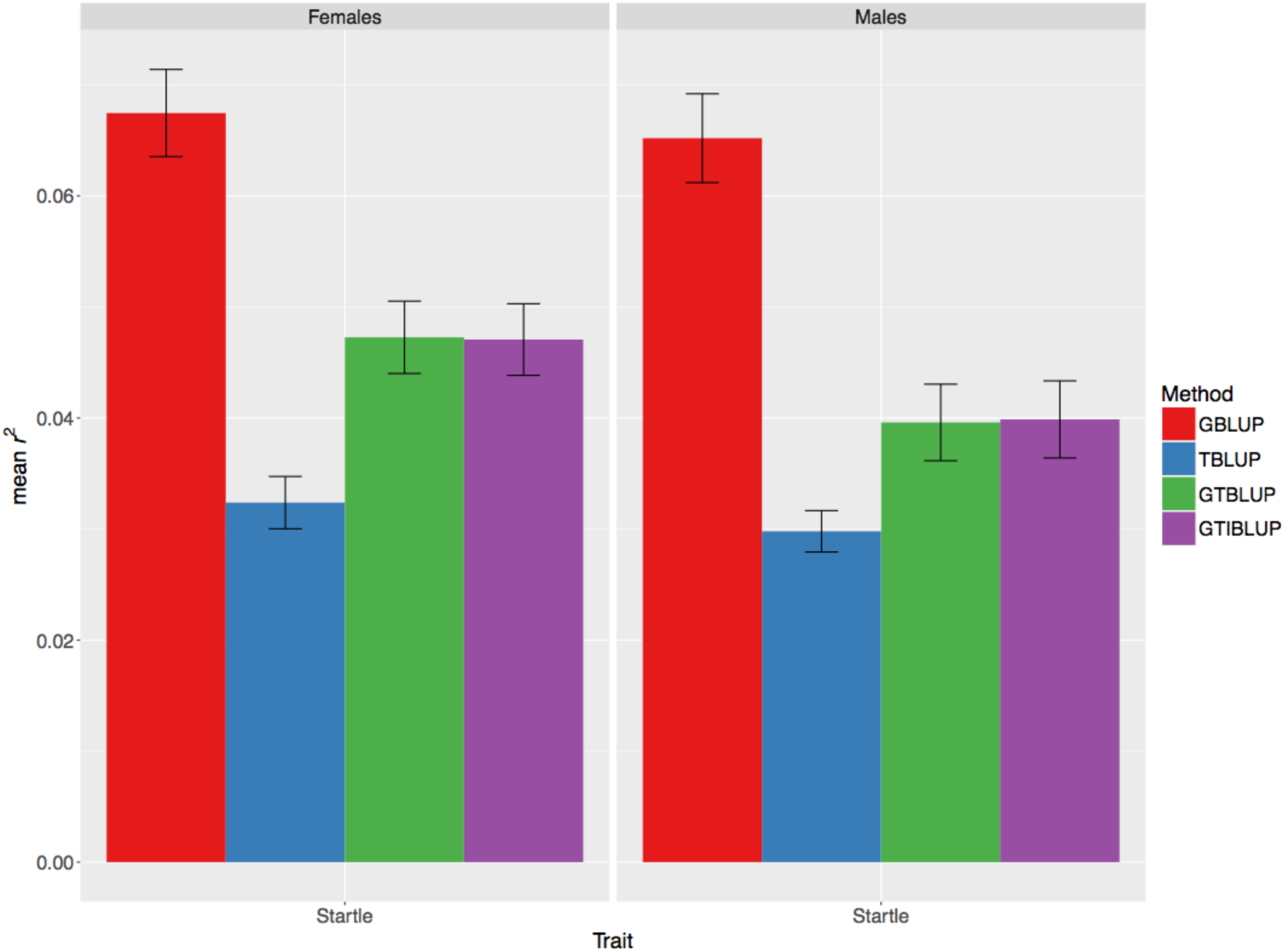
Prediction accuracy for startle response, measured as mean *r*^*2*^ (on the y-axis) in the test set, obtained by GBLUP, TBLUP, GTBLUP, and GTIBLUP. The bars represent the standard error of the mean. The left panel represents females, and the right panel represents males.

**Figure S4.**
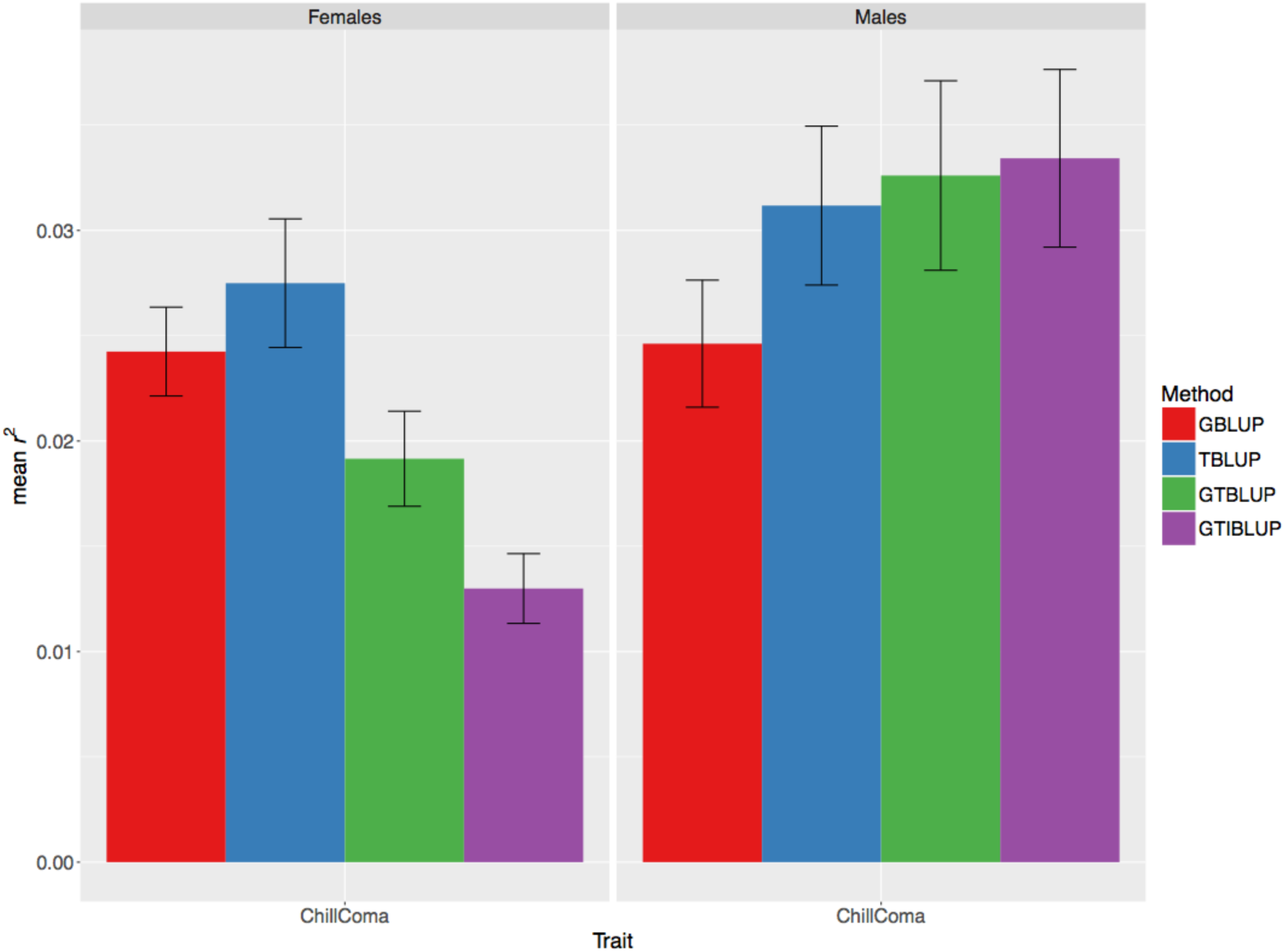
Prediction accuracy for chill coma recovery, measured as mean *r*^*2*^ (on the y-axis) in the test set, obtained by GBLUP, TBLUP, GTBLUP, and GTIBLUP. The bars represent the standard error of the mean. The left panel represents females, and the right panel represents males.

**Figure S5.**
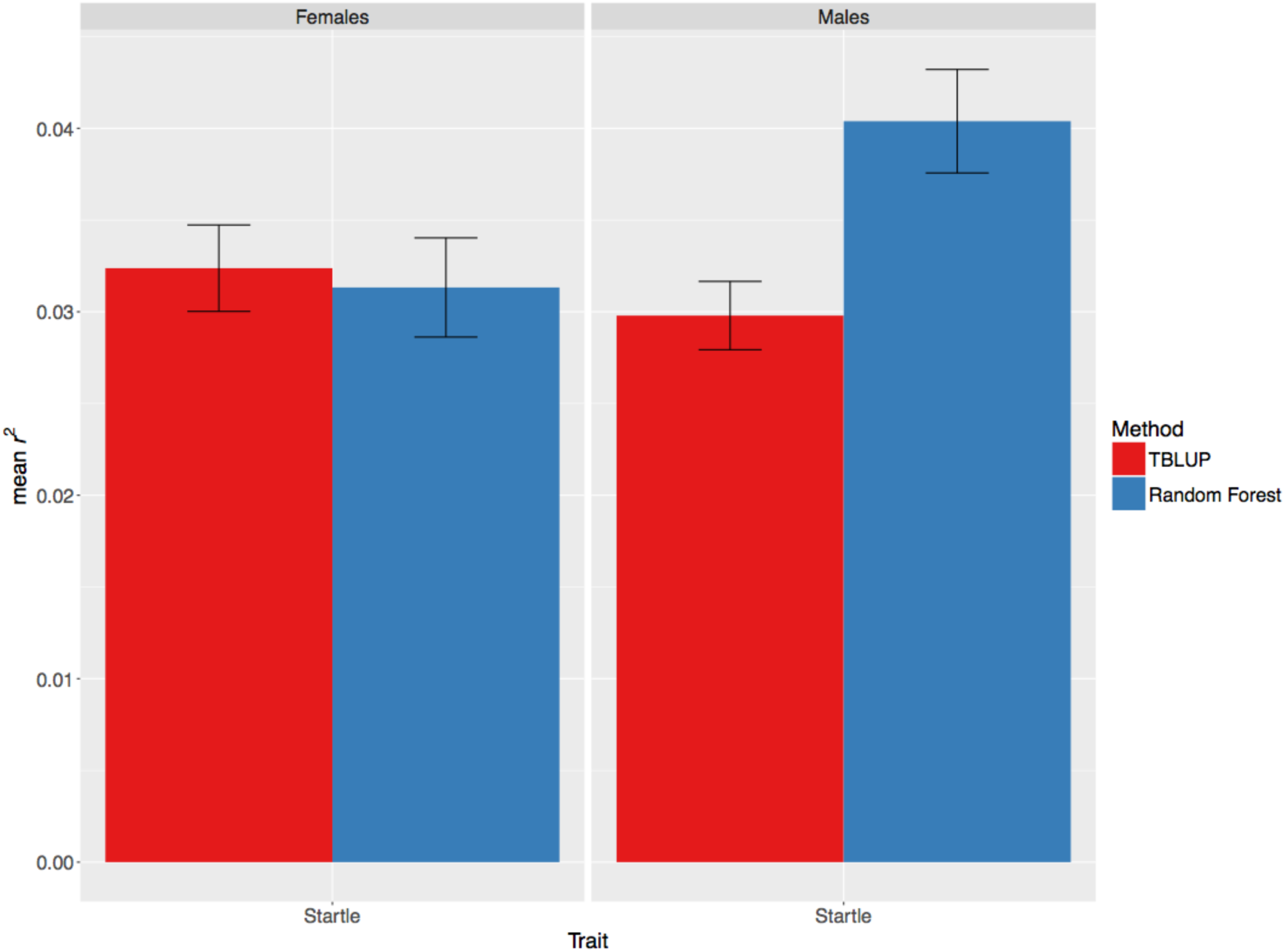
Comparison between the prediction accuracy for startle response, measured as mean *r*^*2*^ (on the y-axis) in the test set, obtained by TBLUP and Random Forest. The bars represent the standard error of the mean. The left panel represents females, and the right panel represents males.

**Figure S6.**
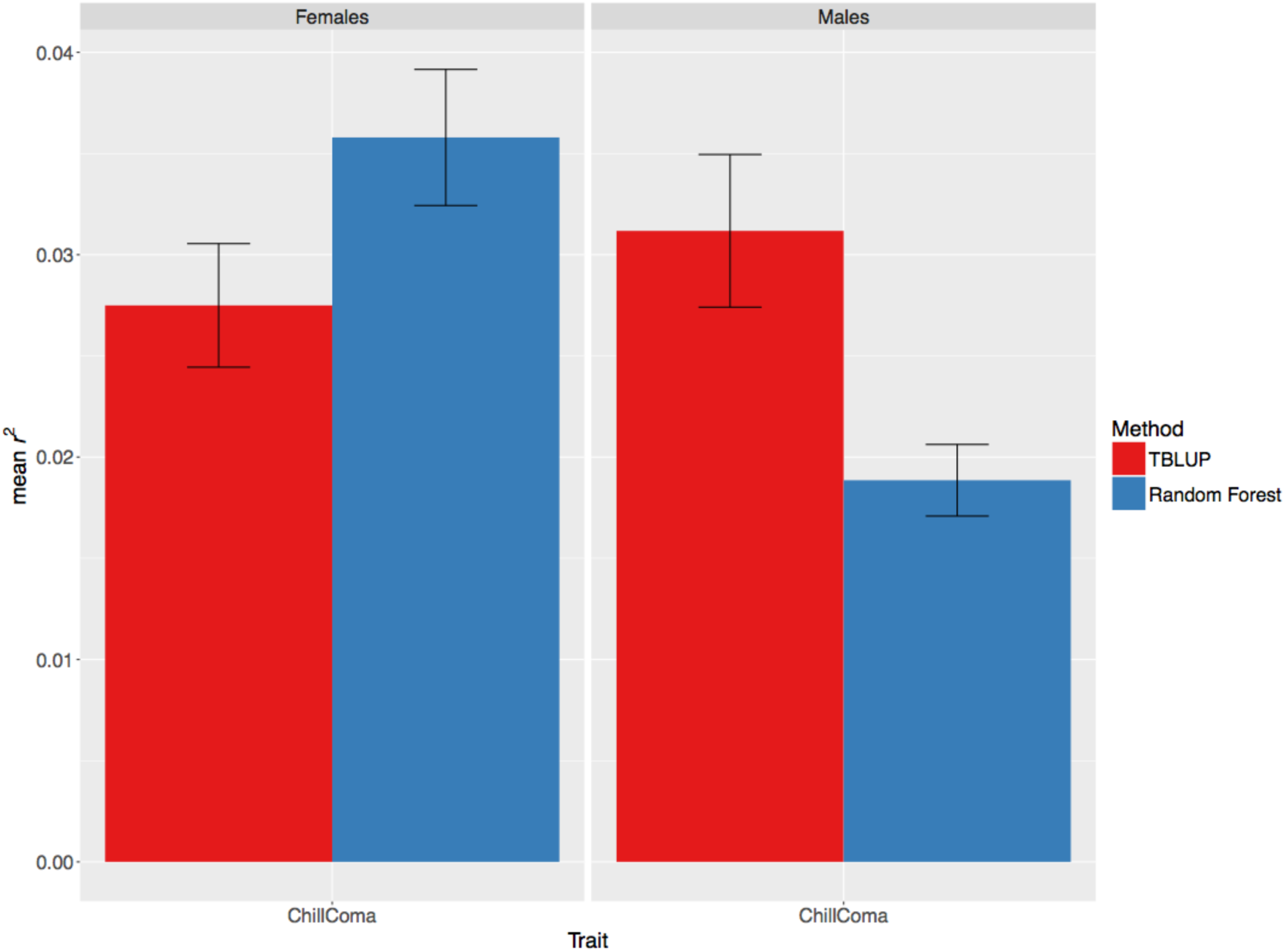
Comparison between the prediction accuracy for chill coma recovery, measured as mean *r*^*2*^ (on the y-axis) in the test set, obtained by TBLUP and Random Forest. The bars represent the standard error of the mean. The left panel represents females, and the right panel represents males.

**Figure S7.**
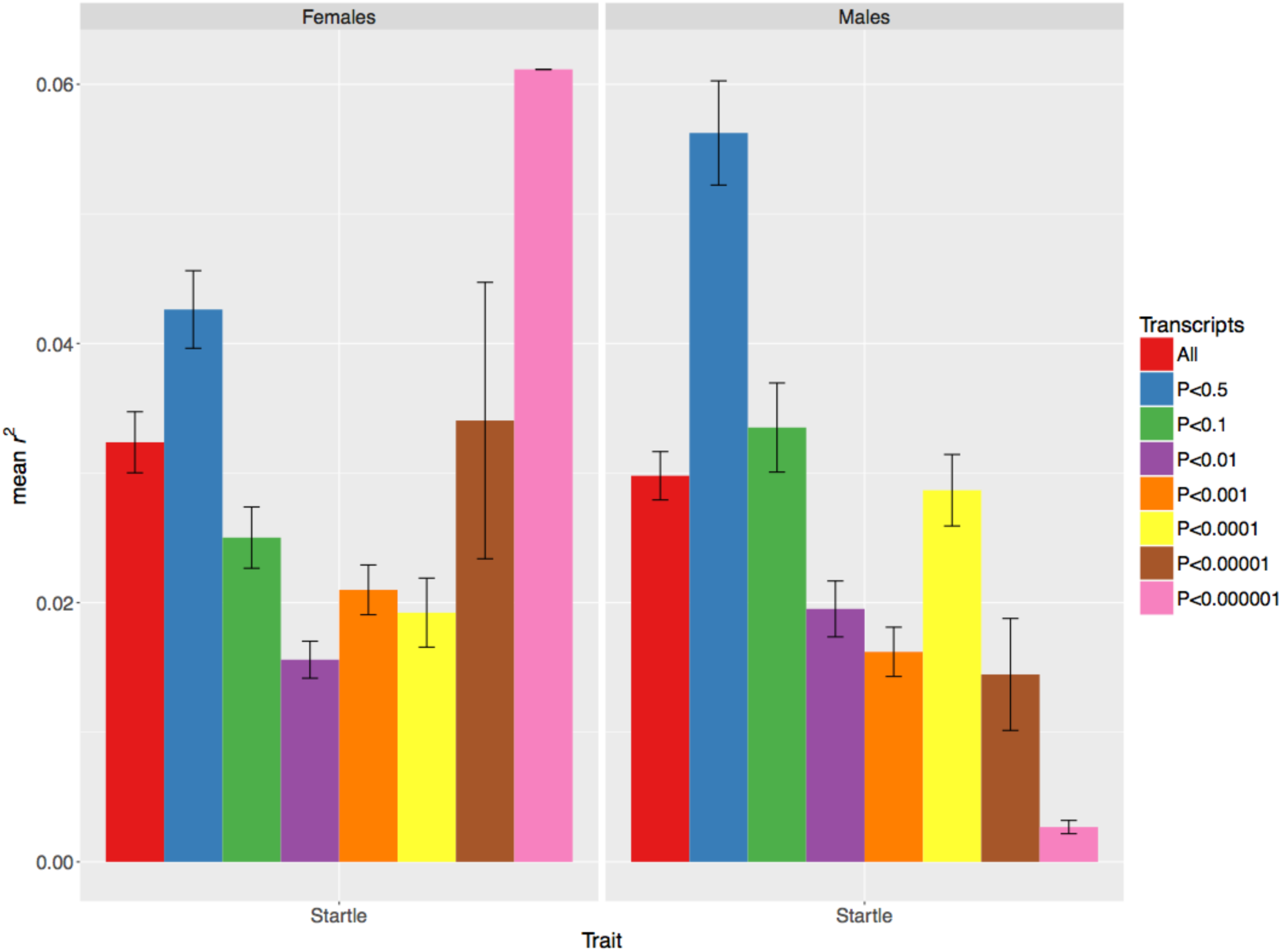
Prediction accuracy for startle response, measured as mean *r*^*2*^ (on the y-axis) in the test set, obtained by the TWAS-TBLUP model at different p-value thresholds for selecting genes. The bars represent the standard error of the mean. The left panel represents females, and the right panel represents males.

**Figure S8.**
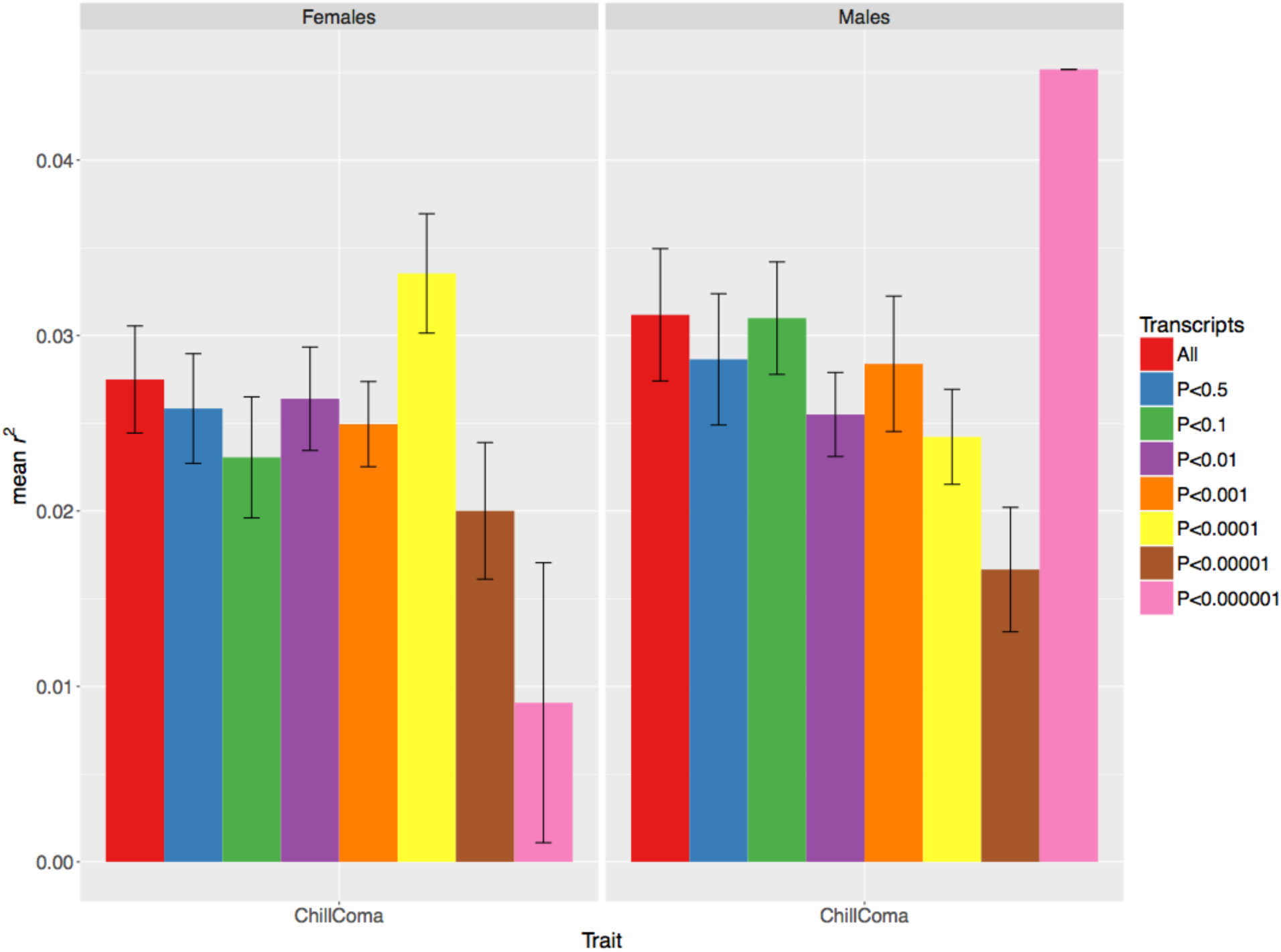
Prediction accuracy for chill coma recovery, measured as mean *r*^*2*^ (on the y-axis) in the test set, obtained by the TWAS-TBLUP model at different p-value thresholds for selecting genes. The bars represent the standard error of the mean. The left panel represents females, and the right panel represents males.

**Figure S9.**
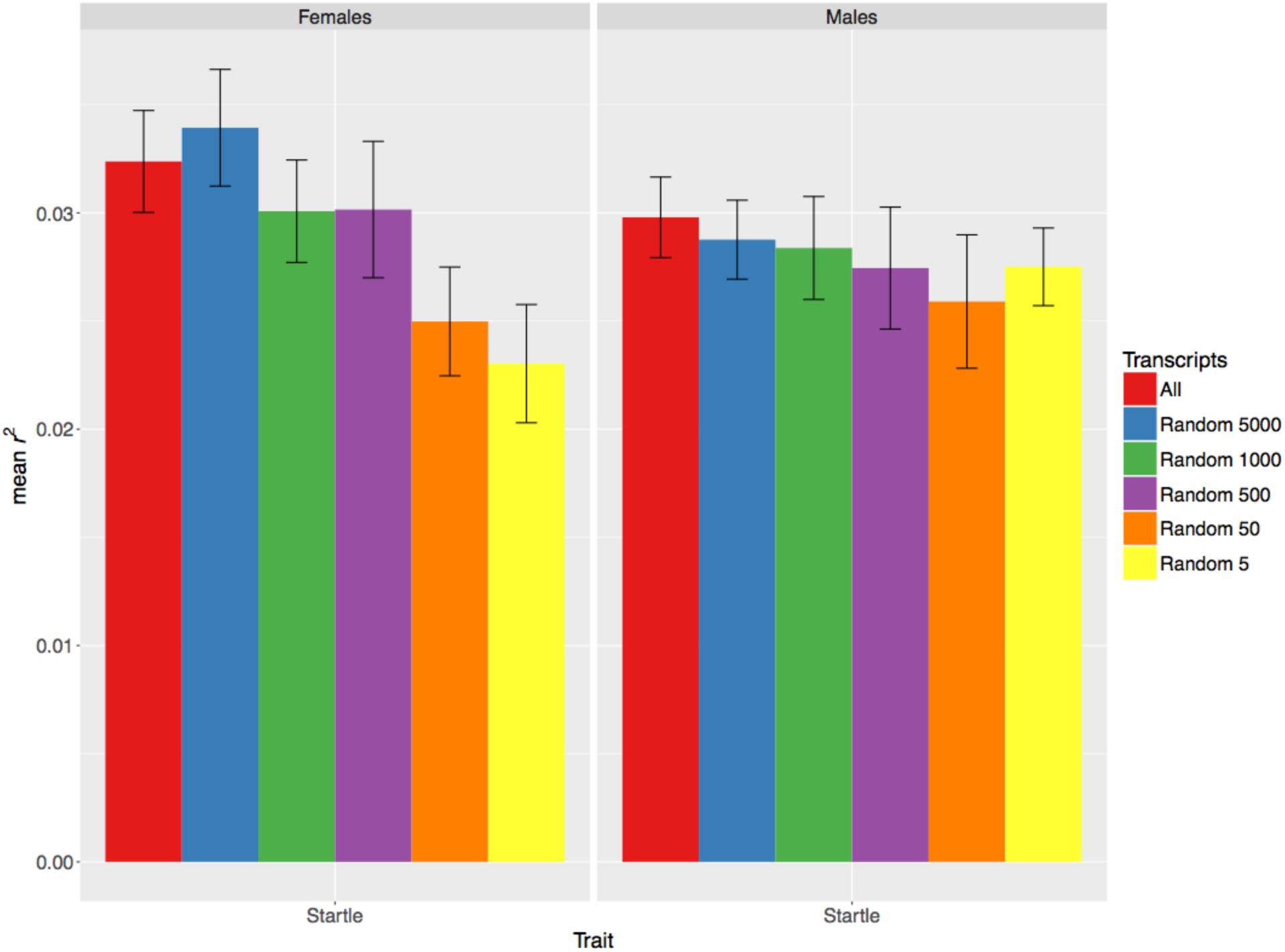
Prediction accuracy for startle response, measured as mean *r*^*2*^ (on the y-axis) in the test set, obtained by using only different numbers of randomly selected genes in the TBLUP model. The bars represent the standard error of the mean. The left panel represents females, and the right panel represents males.

**Figure S10.**
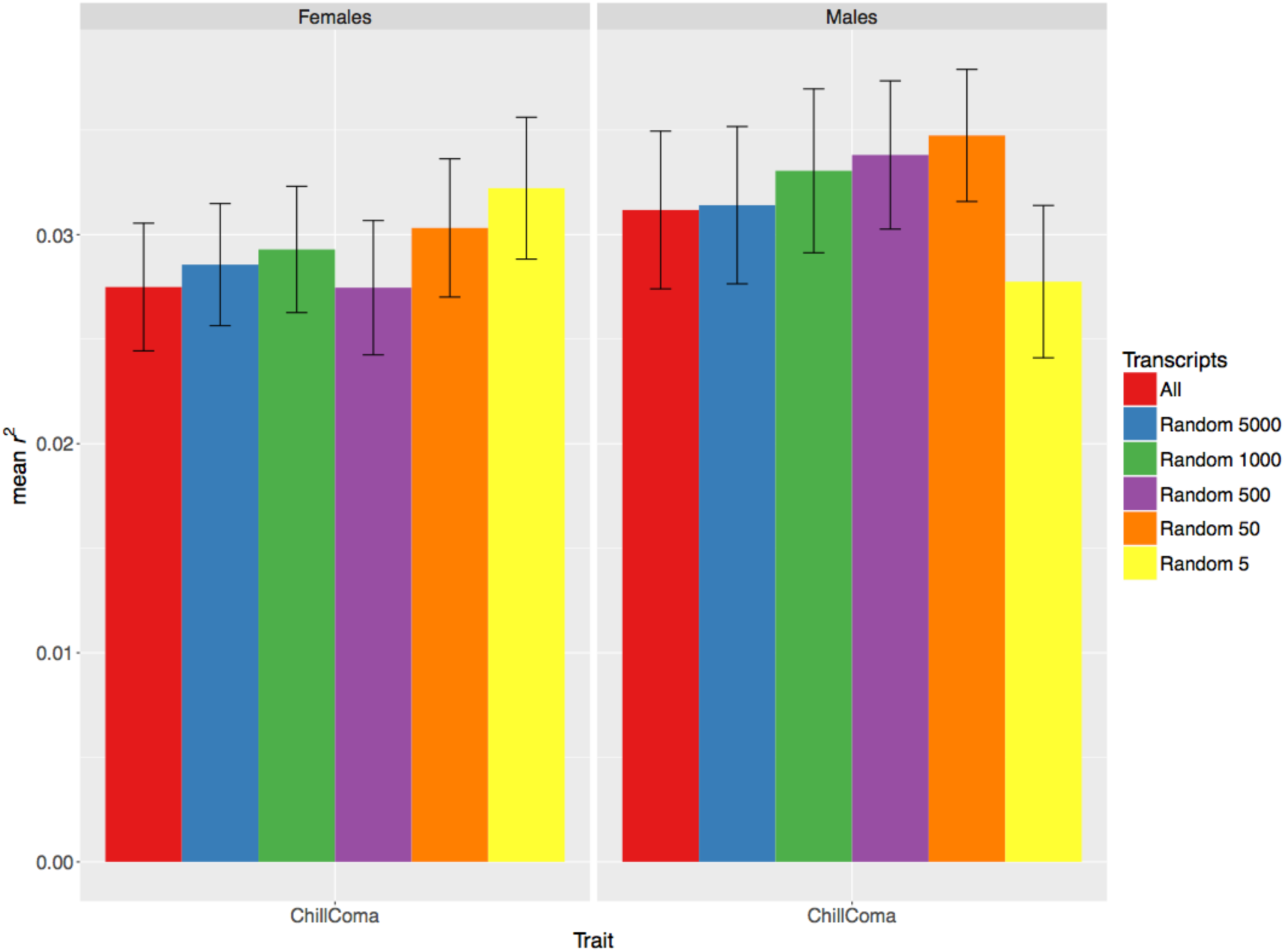
Prediction accuracy for chill coma recovery, measured as mean *r*^*2*^ (on the y-axis) in the test set, obtained by using only different numbers of randomly selected genes in the TBLUP model. The bars represent the standard error of the mean. The left panel represents females, and the right panel represents males.

**Figure S11.**
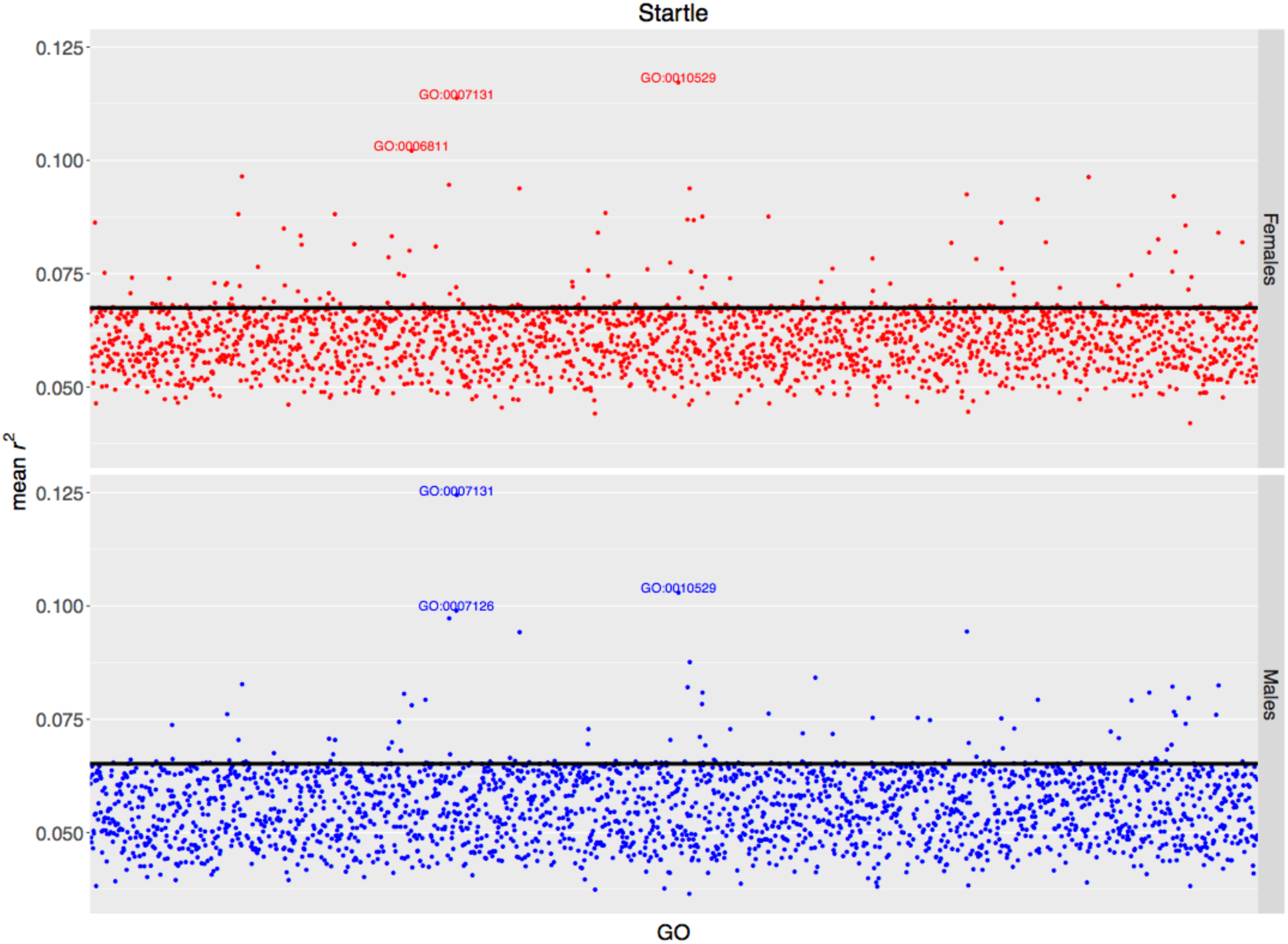
Prediction accuracy for startle response, measured as mean *r*^*2*^ (on the y-axis) in the test set, obtained by the GO-GBLUP model. Each point represents the prediction accuracy achieved by a specific GO term; the top 3 most predictive GO terms are spelled out. The black horizontal line represents the accuracy of the baseline GBLUP model. The upper panel represents females, and the lower panel represents males.

**Figure S12.**
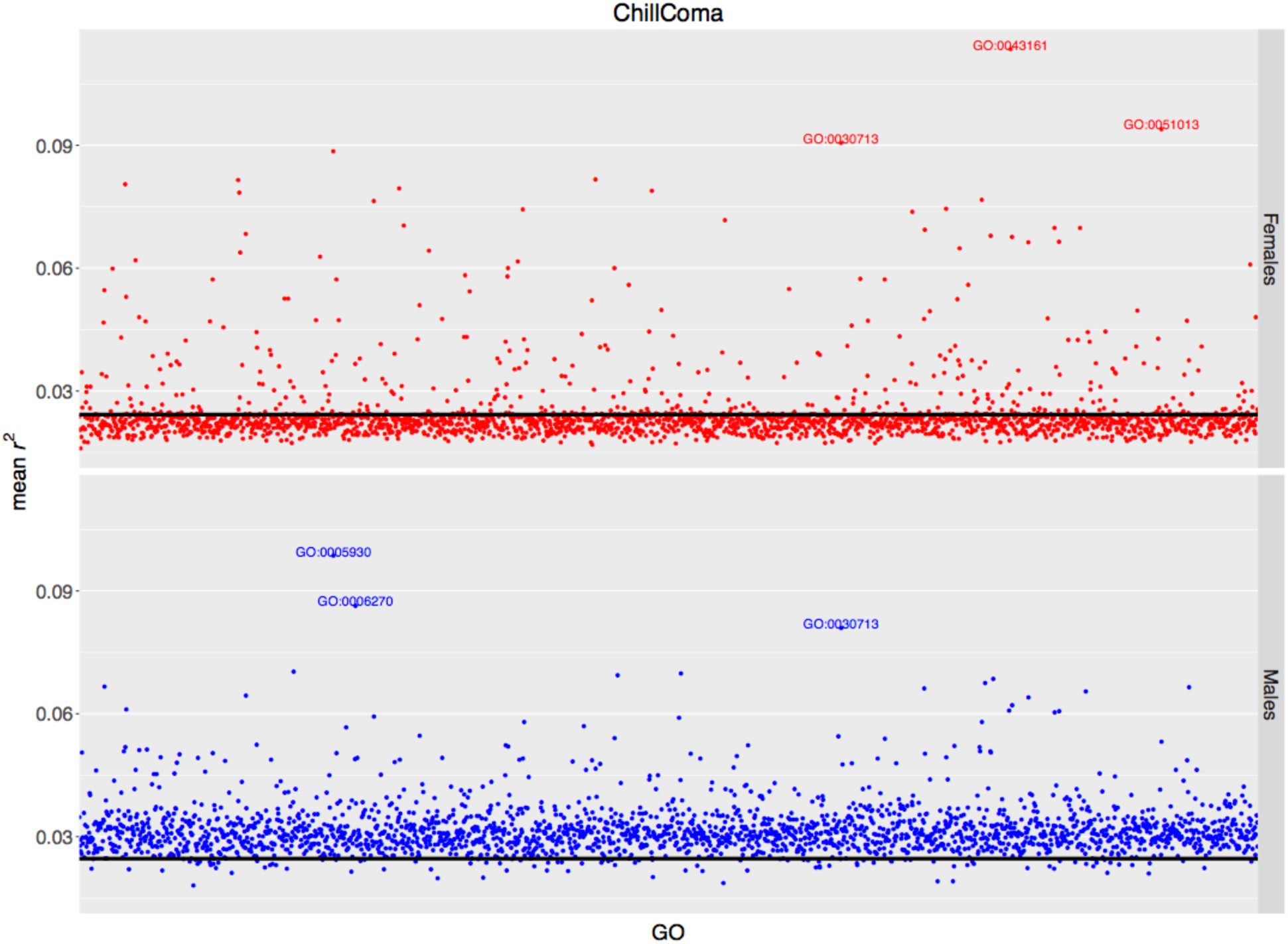
Prediction accuracy for chill coma recovery, measured as mean *r*^*2*^ (on the y-axis) in the test set, obtained by the GO-GBLUP model. Each point represents the prediction accuracy achieved by a specific GO term; the top 3 most predictive GO terms are spelled out. The black horizontal line represents the accuracy of the baseline GBLUP model. The upper panel represents females, and the lower panel represents males.

**Figure S13.**
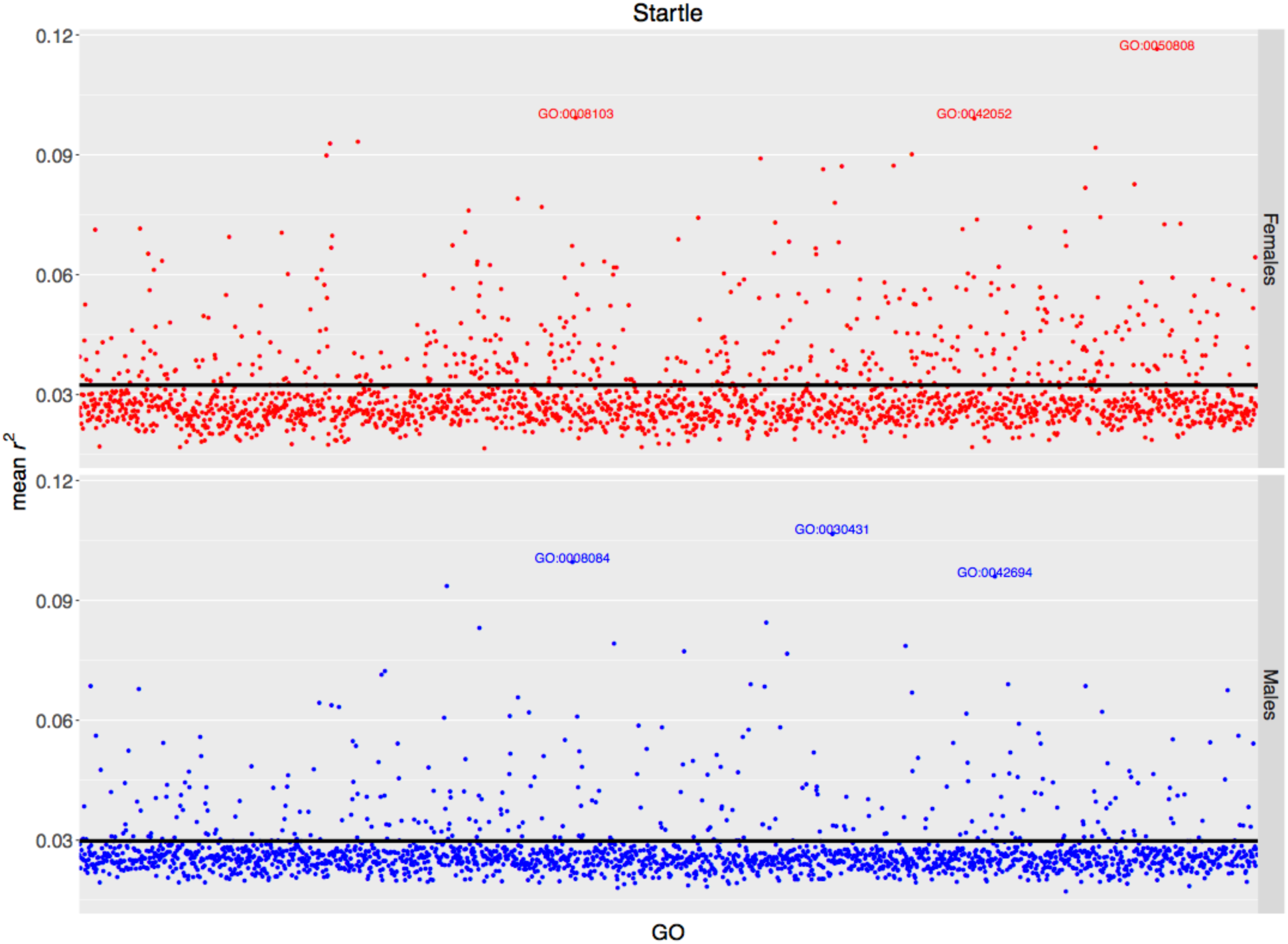
Prediction accuracy for startle response, measured as mean *r*^*2*^ (on the y-axis) in the test set, obtained by the GO-TBLUP model. Each point represents the prediction accuracy achieved by a specific GO term; the top 3 most predictive GO terms are spelled out. The black horizontal line represents the accuracy of the baseline TBLUP model. The upper panel represents females, and the lower panel represents males.

**Figure S14.**
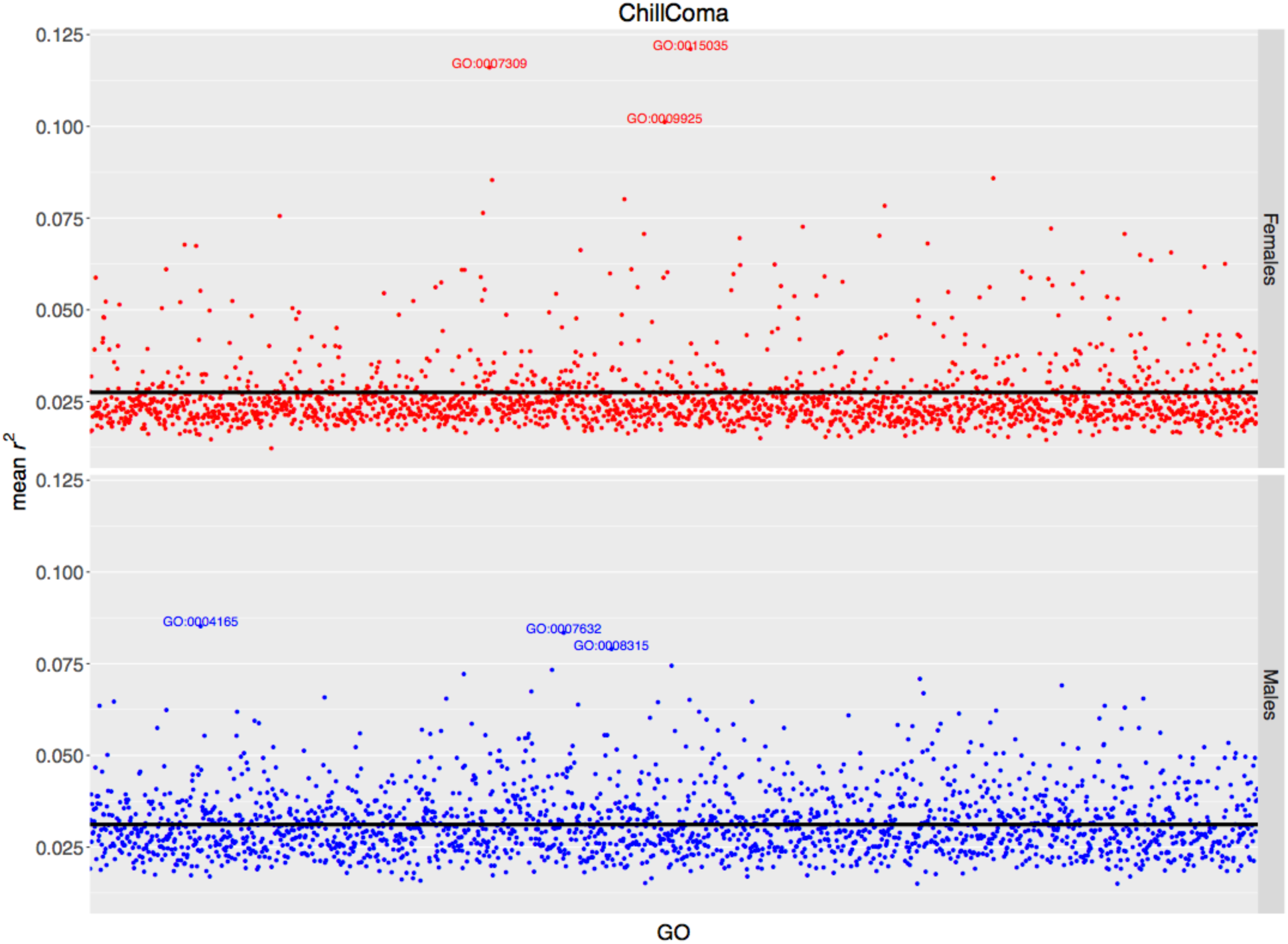
Prediction accuracy for chill coma recovery, measured as mean *r*^*2*^ (on the y-axis) in the test set, obtained by the GO-TBLUP model. Each point represents the prediction accuracy achieved by a specific GO term; the top 3 most predictive GO terms are spelled out. The black horizontal line represents the accuracy of the baseline TBLUP model. The upper panel represents females, and the lower panel represents males.

**Figure S15.**
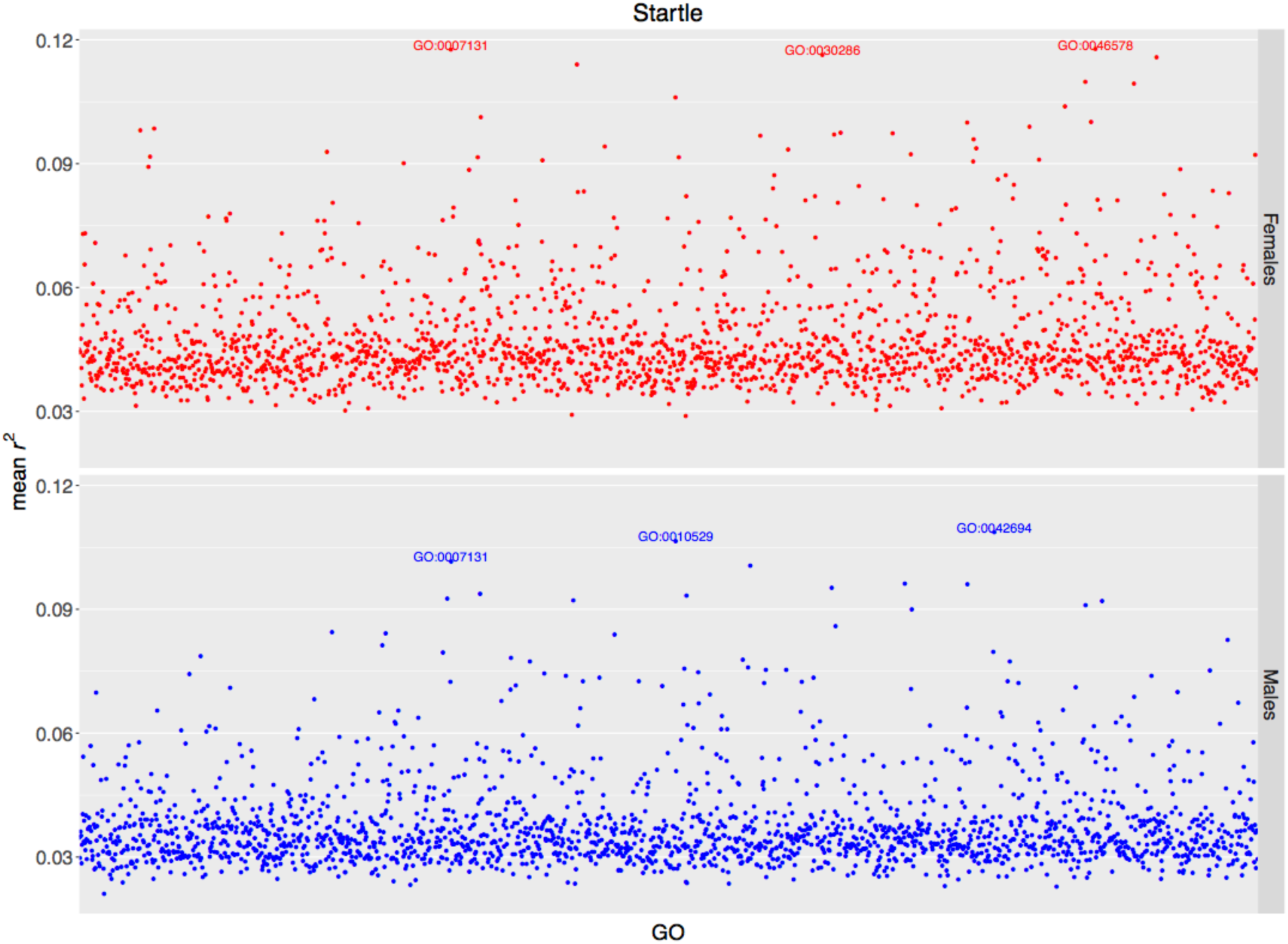
Prediction accuracy for startle response, measured as mean *r*^*2*^ (on the y-axis) in the test set, obtained by the GO-GTBLUP model. Each point represents the prediction accuracy achieved by a specific GO term; the top 3 most predictive GO terms are spelled out. The upper panel represents females, and the lower panel represents males.

**Figure S16.**
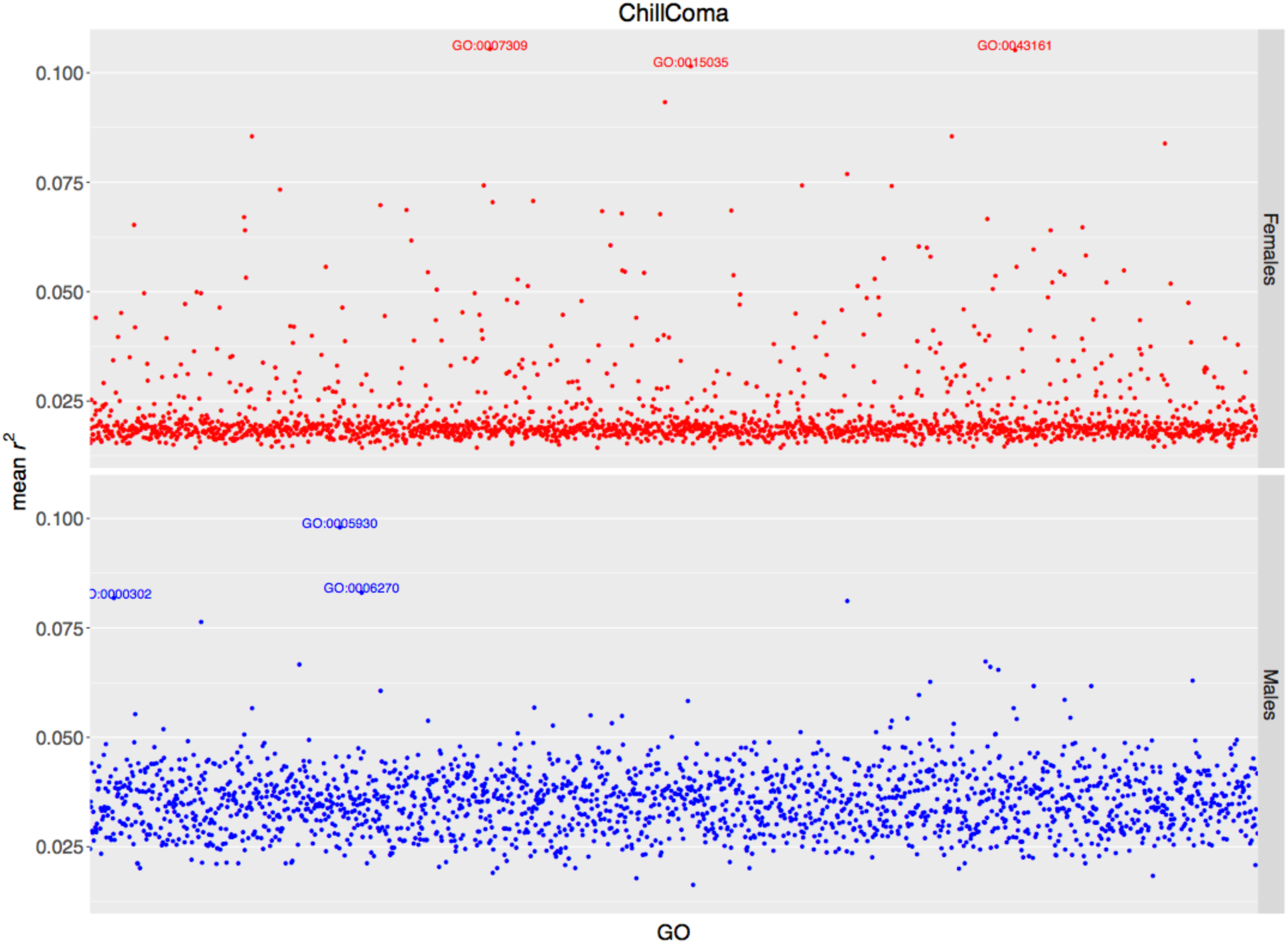
Prediction accuracy for chill coma recovery, measured as mean *r*^*2*^ (on the y-axis) in the test set, obtained by the GO-GTBLUP model. Each point represents the prediction accuracy achieved by a specific GO term; the top 3 most predictive GO terms are spelled out. The upper panel represents females, and the lower panel represents males.

